# Formation of extracellular vesicles depends on mechanical feedback of the cortex and the glycocalyx

**DOI:** 10.1101/2025.06.14.659723

**Authors:** Ke Xiao, Padmini Rangamani

**Author notes:** To whom correspondence must be addressed.

## Abstract

Cell-secreted extracellular vesicles (EVs) play a pivotal role in local and distant cell-to-cell communication by delivering specific cargoes to other cells or to the extracellular space. In many cells, the glycocalyx, a thick sugar-rich layer at the cell surface, and the membrane-cortex attachment are crucially linked to the formation of EVs, yet it is unclear what determines the successful formation of EVs when multiple physical factors are involved. In this work, we developed a model for glycocalyx-membrane-cortex composite to investigate the effects of glycocalyx and membrane-cortex adhesion on the formation of EVs by combining polymer physics-based theory and Helfrich membrane theory. By performing linear stability analysis, we show that modulating the mechanical feedback among the glycocalyx, membrane-cortex attachment, and membrane curvature can give rise to two types of instabilities: a conserved Turing-type instability and a Cahn-Hilliard-type instability. Furthermore, using an equilibrium model, we identified two critical conditions for EV formation: an initial detachment of the membrane from the underlying cortex and then a sufficient driving force to induce membrane deformation for successful EV formation. We further demonstrated that there exists an optimal glycocalyx coating area at which the formation of EVs is most favorable. Finally, we use our model to predict that a heterogeneous size distribution of EVs can be generated through the regulation of glycocalyx properties, shedding insight into how EVs of different radii may be generated.

**Significance Statement:** Extracellular vesicles (EVs) are important for cell biology because they facilitate active communication between cells. Understanding the governing factors that control the formation of EVs is crucial to many cellular processes ranging from tumor progression and metastasis evolution to the disposal of unwanted biomolecules. However, whether EV secretion is a consequence of the glycocalyx and the role of membrane-cortex adhesion in the formation of EVs are still elusive. To address these issues, here we develop a biophysical model for EV formation that couples the presence of glycocalyx and membrane-cortex adhesion. We find that the glycocalyx-membrane-cortex composite system exhibits two types of instabilities utilizing stability analysis – a Turing instability and a Cahn-Hilliard type instability. Based on our proposed equilibrium model, we identified that for the initiation of membrane detachment and the formation of EVs each need to meet a critical threshold. In addition, our model predicts that the formation of EVs is most favorable when an optimal glycocalyx coating area reaches and a heterogeneous distribution EV sizes can be produced by regulating glycocalyx properties.

## Introduction

Extracellular vesicles (EVs) are lipid bilayer-enclosed particles ubiquitously secreted by all cells (*1*– *7*) into the extracellular environment under a range of conditions, from normal and stress states to physiological and pathological states (*1–3, 8, 9*). Various types of cellular components and biomolecules, such as signaling proteins, lipids, nucleic acids (mRNAs, miRNAs, DNA), metabolites, and even pharmacological compounds (*1–3, 10–12*), are shuttled to the extracellular space through many EV subtypes, i.e., exosomes, microvesicles, apoptotic bodies, as well as migrasomes and large oncosomes (*1–4*). The ability to transfer biomolecular cargo into target cells enables EVs to play a pivotal role in biological processes such as local and long-distance inter-cellular communication and the disposal of unwanted materials (*1–4*). In addition, a plethora of studies have reported that EVs also contribute to numerous pathological diseases such as cancer, heart disease, infectious diseases, and immunological and neurological diseases (Alzheimer’s and Parkinson’s) (*7, 13–17*). For example, many cell stress-inducing conditions, such as inflammation, proapoptotic conditions, and hyperbaric pressure, can lead to an increase in the formation of membrane vesicles (*18*). Many different cell types including cancer cells, platelets, and endothelial cells have been known to release EVs and EVs have been implicated in many different physiological and pathological conditions (*19–21*). Based on these definitions and classifications, these extra-cellular vesicles (EVs) generated by direct outward budding or blebbing of specific sites of the plasma membrane are generally referred to as ectosomes or microvesicles (MVs) (see Fig. 1(a)). Hence, EVs are fragments of the plasma membrane secreted in response to activating stimuli such as oxidative stress, cytokines, and mechanical forces. In general, EVs are heterogeneous spherical phospholipid bilayer structures with sizes spanning from approximately tens to thousands of nanometers (*22, 23*). The categorization of heterogeneous EVs population is summarized in Table 1. However, the biophysical mechanisms of generating EVs across a spectrum of sizes, along with the probability of their size occurrences, remain poorly understood.

**Table 1.** Types of extracellular vesicles.

| Name | Category | Size range | Origin | Type of cells | References |
| --- | --- | --- | --- | --- | --- |
| Exosomes | Small EV | ~ 30–120 nm | Inward budding of endosomal compartments | Normal cells and cancerous cells | (1–3, 5, 6, 24, 25) |
| Small ectosomes | Small EV | ~ 30–150 nm | Direct outward budding of the plasma membrane | Healthy cells, perivascular dendritic cells, T cells, and cancer cells | (1, 3, 5, 6, 26, 27) |
| Microvesicles | Small to large EV | ~ 150–1000 nm | Direct outward budding of the plasma membrane | Epithelial cells, normal cells, and cancerous cells | (1–3, 5, 6, 28, 29) |
| Apoptotic EVs | Small to large EV | ~ 100–5000 nm | Fragments released by dying cells in the late stage of apoptosis | Dying cells | (1–3, 5, 6, 30, 31) |
| Migrasomes | Large EV | ~ 500–3000 nm | Retraction fibers | Migrating cells | (1, 3, 5, 6, 32–34) |
| Oncosomes | Large EV | ~ 1–10 μm | The shedding of non-apoptotic plasma membrane blebbing | Tumor cells | (1–3, 5, 6, 35, 36) |

**Figure 1.**
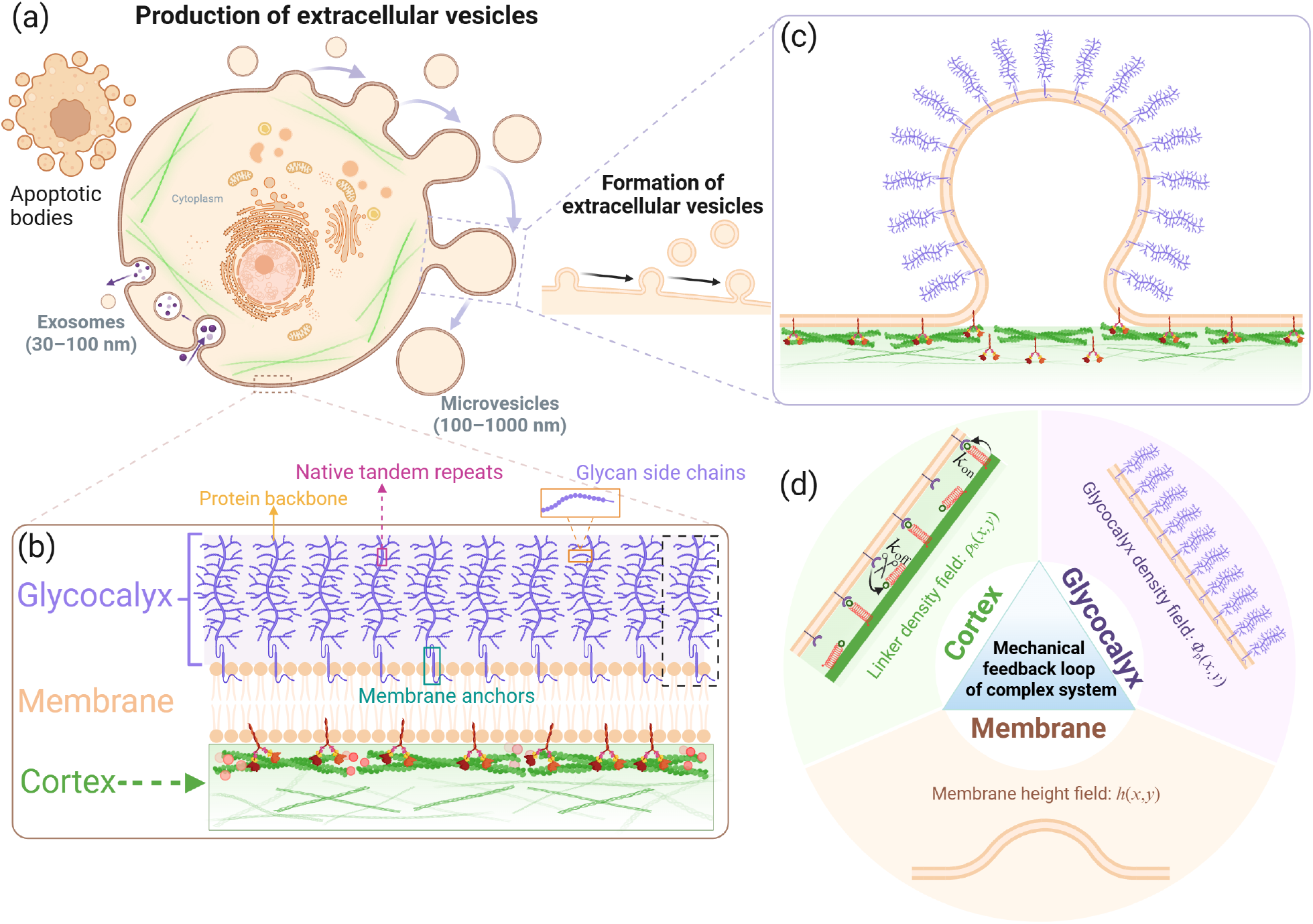
Schematic representation of the factors involved in the formation of EVs. (a) Production and release of EVs into the extracellular space. (b) Schematic representation of three main factors – glycocalyx, membrane, and cortex – considered in this study for the formation of EVs. (c) An enlarged view of an EV detached from the underlying cortex driven by the glycocalyx. (d) Sketch of the glycocalyx-membrane-cortex composite system. In this work, we explore the feedback between these three elements using biophysical modeling. Panels are prepared using Biorender.com.

Many experimental studies have characterized different factors that affect the formation of membrane geometries and we briefly summarize them here grouped by protein effects, membrane effects, or cytoskeletal effects (*37, 38*). One of the predominant views has been that the mechanisms for the generation of curvature are attributed to protein insertion, scaffolding, and crowding (*37–44*). However, a majority of the modeling studies are focused on inward membrane bending such as endocytosis (*45–48*). Instead, a number of membrane morphologies (EVs, cellular blebs, spherical protrusions, tubular protrusions, microvilli, etc.) are generated by bending the membrane outward, and these structures play a crucial role in a spectrum of biological functions and processes in many cells (*49*). Additionally, during membrane shape remodeling, numerous biomacromolecules, membranes, and cytoskeletal structures inside the cell are involved. For example, the glycocalyx (see Fig. 1(b)), a thick layer of heavily glycosylated transmembrane biomacromolecule proteins concentrated on most cell surfaces in a complex brush structure (*50–53*), has been reported to regulate the formation of spherical vesicles, pearls, and tubes along the plasma membrane. As a result, another recently acknowledged membrane curvature generating mechanism is that in order to reduce the entropy cost of the system, the glycocalyx on the outer leaflet tends to drive membrane bending so as to provide more space and additional orientational degrees of freedom for glycopolymer chains (*38, 53–55*). In addition, interactions between the membrane and the cortex also control membrane shape changes; for instance, the formation of cellular blebs requires the detachment of the membrane from the underlying cortex (*56–63*) and cell membrane protrusion initiation is triggered by actin-membrane detachment (*64–67*). The cortex is attached to the membrane through a large number of linker molecules (*49, 68, 69*) such as talin (*63, 70*) and the ezrin, radixin, and moesin (ERM) family of proteins (*71–73*). The disruption of links between the cortical cytoskeleton and the plasma membrane is a required condition for successful membrane detachment (see Fig. 1(c)). A fundamental challenge in predicting EV formation regulated by glycocalyx and membrane-cortex adhesion stems is to understand the energetic contributions of the glycocalyx, membrane-cortex adhesion, and associated factors, such as glycocalyx area coverage, line tension created by multiple phases on the membrane, spontaneous curvature owing to asymmetries in the bilayer, and membrane tension (*45, 46, 74, 75*).

Since EVs and highly curved membrane shapes are essential to diverse cellular functions, the formation of membrane morphologies has been extensively studied by experiments and theory. Experimentally, it has been shown that polypeptide and a class of large, heavily glycosylated proteins called mucins are frequently accumulated at high densities on protrusive membrane structures such as epithelial microvilli (*76–81*), cilia (*82*), and filopodia (*83, 84*). More recently, Shurer et al. (*53*) reported that bulky brush-like glycopolymers are sufficient to induce a variety of curved membrane features, including spherical-shaped membranes (referred to as blebs), tubes, and unduloids, in a density-dependent manner. Theoretical studies have used mean-field theory, analytical methods (*85–88*), scaling theory (*89*), phase field models (*90, 91*), and Monte Carlo simulations (*92, 93*), to show that anchored polymers on membrane give rise to changes in the shape, elastic properties, and spontaneous curvature of the plasma membrane. However, the membrane response to a thick layer of glycocalyx has not been well investigated. Recently, we developed a general energetic framework that couples the mechanics of the glycocalyx with the mechanics of the lipid bilayer to investigate how the glycocalyx can generate spherical vesicles and tubular structures (*74, 75*). Meanwhile, there are plenty of studies examining the role of the cortex in bleb formation by combining experimental observations and continuum mechanics models (*57–60, 94, 95*). Also, a biophysical model is recently developed to account for the interplay between cortical adhesion and membrane bending in the regulation of microparticle formation (*96*). Despite the fact that EVs have become the focus of rising interest and it is well known that they are formed by outward membrane bending, the balance of forces required for EV formation is not yet well understood.

To address this gap, in this work, we develop a theoretical model to interpret the formation of EVs based on polymer physics-based theory and Helfrich membrane theory, which captures the combined effects of glycocalyx, membrane-cortex adhesion, glycocalyx coating area, line tension, spontaneous curvature, and membrane tension. Based on this framework, we systematically investigated the influence of different factors on the formation of EVs (Fig. 1(d)). Specifically, we begin our exploration by providing a linear stability analysis of the glycocalyx-membrane-cortex complex system (Sec. 1). Next, we propose an equilibrium model to study the role of each factor in the formation of EVs (Sec. 2). Our model predicts that (i) the complex system becomes unstable under the conditions of depletion of the adhesive protein linkers and increasing the density and length of glycopolymers; two types of instabilities can emerge by tuning the control parameters in the unstable regime. (ii) A critical condition is required for the onset of membrane detachment from the cortex, and the formation of a complete EV requires a threshold grafting density. (iii) There is an optimal glycocalyx coating area for the formation of EVs and a wide range of EV size can be generated by regulating glycocalyx properties. This work provides a comprehensive model for EV formation by a detailed representation of the plasma membrane going beyond the lipid bilayer.

## Model Development and Methods

To investigate the influence of glycocalyx and membrane-cortex attachment on EV formation, we first construct a minimal dynamical model (see Sec. 1) that takes into account both glycopolymer-membrane and membrane-cortex interactions and study the stability of this complex system. Then, we develop a biophysical model (see Sec. 2) to investigate the effects of glycocalyx and membrane-cortex adhesion on membrane shape remodeling (EV formation). We consider a system comprising a cell membrane coated with glycocalyx on the outside of the surface and attached to a thin layer of cortex through protein linkers, as shown in Fig. 1. The total free energy of the system is given by the sum of the following three contributions, i.e.,

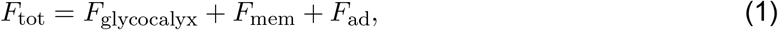

where the first contribution is the energy associated with the glycocalyx, the second contribution is the Canham-Helfrich free energy due to bending the membrane (*97, 98*), and the last term is the membrane-cortex adhesion energy, which stems from the attachment of the membrane to the cortex.

The first energy term related to the glycocalyx is described separately in Sec. 1A and Sec. 2B. The second energy term on the right-hand side of Eq. (1) can be given by the Canham-Helfrich Hamiltonian (*97, 98*) as

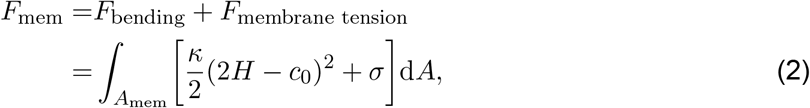

where *κ* is the bending modulus, *H* = (*c*_1_ +*c*_2_)*/*2 and *c*_0_ are the mean and spontaneous curvatures, *c*_1_ and *c*_2_ are the two principal curvatures, *σ* is the membrane surface tension. In Eq. (2), the first term accounts for the bending energy, which arises from the membrane resistance to the out-of-plane deformation, whereas the last term describes the tension energy due to membrane stretching.

Since membrane-cortex adhesion couples to membrane deformations, we also consider the effect of the tethering of the membrane to the cell cortex. To quantitatively capture this effect on EV formation, we assume that the attachment between the cell membrane and the actin cortex through linker proteins and the linker proteins (i.e., ezrin, radixin, and moesin) behave as springs arranged in parallel. Thus, the membrane-cortex adhesion energy cost due to tethering of the membrane to the cortex can be treated as quadratic with respect to the membrane-cortex linker deformation and is given by a harmonic potential of the form (*56, 57*)

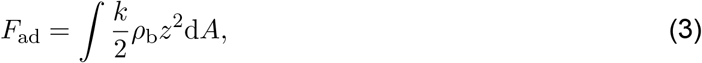

where *k* is the effective elastic stiffness (spring constant) of each membrane-cortex linker, *ρ*_b_ is the local density of bound spring-like linkers, and *z* is the stretching length of linkers.

### 1 Onset of glycopolymer phase separation and initiation of EV formation

To explore whether the glycocalyx undergoes phase separation on the cell membrane and initiates the formation of EV when the membrane is attached to the cortex, we first propose a free energy functional to derive the kinetic equations that govern the dynamics of the complex system, and then perform linear stability analysis to demonstrate the instability of the system under certain conditions. Hence, we employ the Flory-Huggins theory here to describe the energy related to the glycocalyx on the membrane.

#### A. Free-energy functional

To study the stability of the glycocalyx-membrane-cortex system, we characterize this system by three state variables: [*φ*_p_(***r***), *ρ*_b_(***r***), *h*(***r***)], where *φ*_p_(***r***) represents the area fraction of glycopolymers on the membrane, *ρ*_b_(***r***) denotes the molecular linker density attached to the membrane, and *h*(***r***) is the normal displacement (height) of the membrane above a reference plane parametrized by ***r*** = (*x, y*). We partition the total free energy functional of the system into three contributions

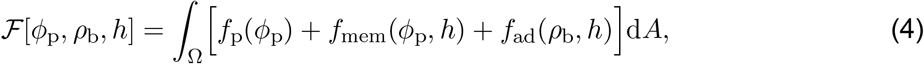

where the first term is the functional for the free energy densities of the glycopolymer on the membrane, the second term represents the Canham-Helfrich free energy (*97, 98*), and the last term denotes the membrane-cortex adhesion energy.

For the functional of the free energy density associated with the glycopolymers on the membrane, we adopt the standard Flory-Huggins model (*99, 100*) to account for the entropy of mixing and the interaction of glycopolymers:

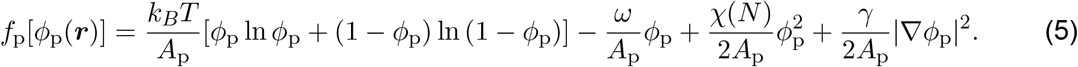

In this equation, *A*_p_ is the surface area occupied by a single grafted polymer on the membrane, *ω* stands for the chemical potential of the polymer binding, *χ* is a direct interaction coefficient between glycopolymers, and *γ* sets the energetic penalty for composition gradients. Note that the length of glycopolymers also affects the inter-polymer interactions here. For simplicity, we assume that *χ* depends on *N* through the relation *χ*(*N*) = (1 + *N*)*χ*_0_, where *N* is the number of monomers in a polymer brush. In this work, we use the number of monomers, *N*, to capture the length of the polymer due to that the thickness of polymer brush is related to the total number of monomers (see Eq. (S12)).

Using the Monge representation for membrane deformation under small gradient approximation (|*∇h*|*≪*1), the free energy density for the elastic deformation of the membrane is given by

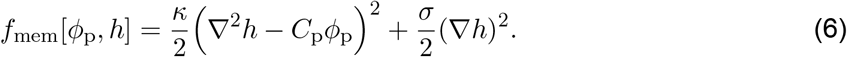

Here, we assume that a linear dependence of the spontaneous curvature on the glycocalyx density *φ*_p_ with a proportionality factor *C*_p_ encodes the curving strength of glycopolymers.

Considering that the local stretching length of linkers is equal to the height of the membrane, i.e., *z* = *h*, the last energy functional density appearing in Eq. (4) can be expressed as

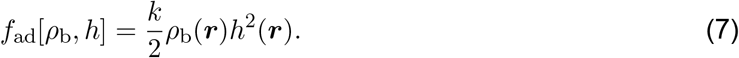

Combining Eqs. (5)-(7), the total free energy functional can finally be written as the sum of these contributions, with

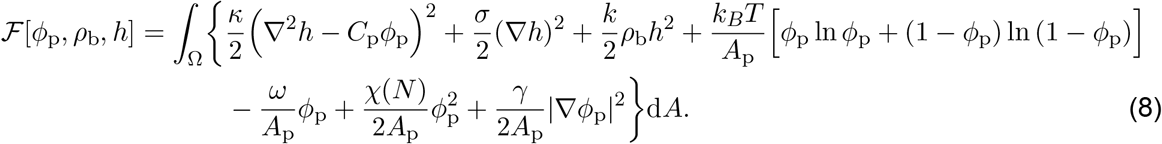

#### B. Coupled dynamics

Assuming that the membrane deforms to reduce the total free energy, the shape-deformation dynamics can be described by an energy relaxation equation. The time-evolution equation for the shape of the membrane is given by

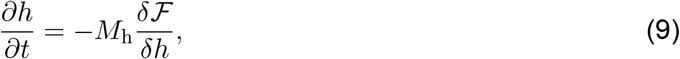

where *M*_h_ is a damping coefficient. Using Eqs. (8) and (9), we find

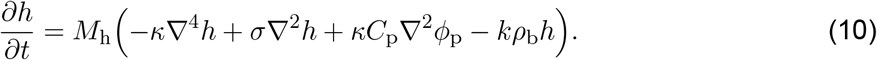

We assume that the number of glycopolymers on the membrane is conserved. The time-evolution equation for *φ*_p_ can be obtained from the continuity equation *∂*_*t*_*φ*_p_ + *∇ ·* **J** = 0 with a current density **J** = *−M*_p_*∇µ*, where *M*_p_ is the mobility coefficient and *µ* = *δ ℱ /δφ*_p_ is the chemical potential. Here *M*_p_ = *D*_diff_ *A*_p_*/*(*k*_*B*_*T*) with *D*_diff_ being the diffusion constant. Performing the functional differentiation one can obtain a Cahn-Hilliard-type dynamical equation for the glycopolymer density as

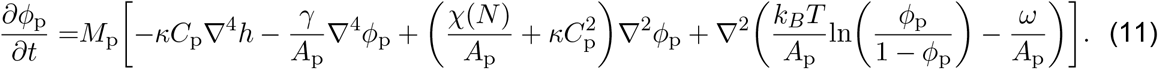

We also assume that the linker protein undergoes turnover between a detached state at a force-dependent rate *k*_off_ and an attached state at a constant rate *k*_on_. Then, the kinetics of tethered linker proteins can be simply described by (*56, 57*)

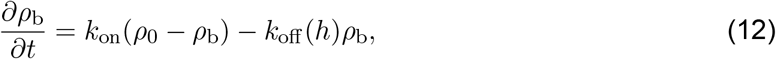

where *ρ*_0_ is the density of available linkers and 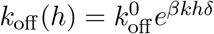 (*101*) with *δ* being a bond length in the nanometric scale (*102*).

#### C. Linear stability analysis

Next we begin from a uniform initial state and look for conditions of instability. We perform a linear stability analysis on the system of equations given in Eqs. (10)-(12), by introducing small perturbations around the spatially homogeneous flat membrane state: *h* = *h*_eq_ + *δh, φ*_p_ = *φ*_p,eq_ + *δφ*_p_, and *ρ*_b_ = *ρ*_b,eq_+*δρ*_b_. Taking the spatial Fourier transform of Eqs. (10)-(12) yields the linearized equations

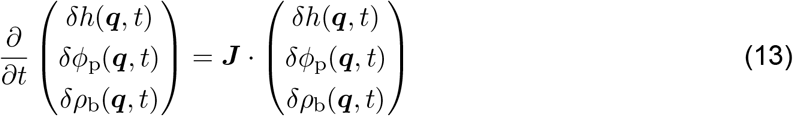

where ***J*** (*q*) = [*J*_*ij*_(*q*)]_3*×*3_ is the Jacobian matrix which reads

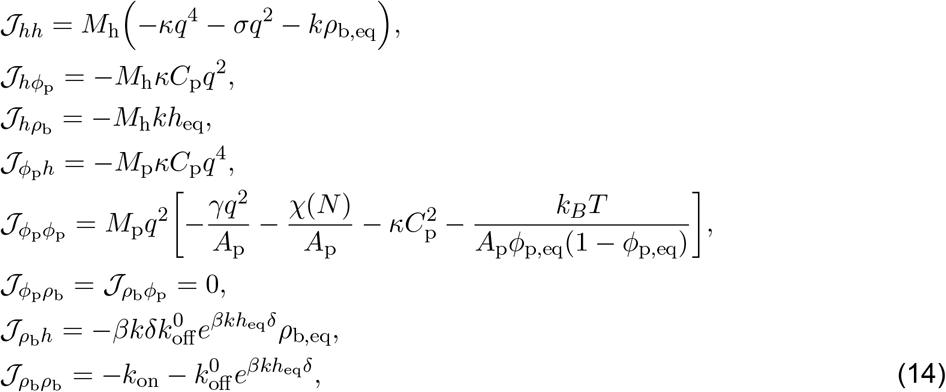

where *q* is the wavenumber. The stability of the glycocalyx coated and cortex attached membrane is governed by the real or complex eigenvalues Λ_*q*_ of the Jacobian associated with the matrix ***J***. The largest eigenvalue of the Jacobian matrix determines the highest growth rate of the perturbations. The dispersion relation Λ(*q*)

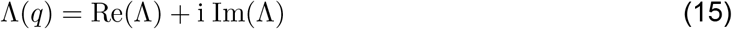

can be determined from the characteristic cubic equation

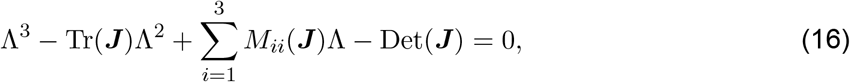

where Tr(***J***) stands for the trace of ***J***, *M*_*ii*_(***J***) for the minors of the diagonal terms of ***J*** and Det(***J***) for the determinant of ***J***. This cubic equation can be solved analytically (*103, 104*) but the resulting expressions are rather complex so we present them here graphically (see the Result Section). The real part Re(Λ) is a rate that characterizes the exponential growth (Re(Λ) *>* 0) or decay (Re(Λ) *<* 0) of the corresponding linear mode. The imaginary part Im(Λ) corresponds to a frequency that can be zero (stationary case) or nonzero (oscillatory case) (*105*). By analyzing the determinant of the rank 3 Jacobian matrix, we can readily identify whether the system is unstable. This is due to the fact that the determinant of ***J*** is equivalent to the product of all its eigenvalues, represented as Det(***J***) = Λ_1_Λ_2_Λ_3_. If Det(***J***) *>* 0, it implies the existence of at least one positive eigenvalue, indicating that the system is unstable.

### 2 Equilibrium description of EV formation

In this section, we consider the scenario in which the glycocalyx aggregates into a stable and area-fixed domain through phase separation. Thus, the energy contribution associated with the glycocalyx is no longer described by the Flory-Huggins theory here. Instead, we adopt the polymer brush-based theory to characterize this energy contribution, which is given by Eq. (19). We assume that this phase-separated domain is flat because of the small deformation approximation. Our aim here is to investigate the conditions under which this process can initiate membrane bending and facilitate EV formation. In the following, we briefly summarize the theoretical model and include additional details in SI.

#### A. Description of the model: Geometry of the EV

To investigate the effects of glycocalyx properties and membrane-cortex adhesion on membrane morphologies, i.e., spherical extensions, we consider a glycocalyx-membrane-cortex system, as shown in Fig. 2. The system we considered consists of a membrane patch that anchored by glycocalyx biopolymers and a thin layer of actin cortex attached beneath the membrane (see Fig. 2). We assume that the patch is much smaller than the average radius of the cell, and this membrane patch with fixed surface area 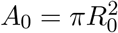 is deformed into a set of spherical caps with radius *R*_s_ due to the presence of glycocalyx, where *R*_0_ is the in-plane radius of the flat membrane (see Fig. 2). The configuration of the membrane system can then be described by a single shape parameter *η* which characterizes the area ratio between a spherical cap and a sphere. In the one-parameter family of configurations of the spherical cap, *η* = 0 corresponds to a completely flat circular bud with radius *R*_0_, while at *η* = 1 the bud is a sphere which implies an EV is formed. The membrane domain takes the shapes of spherical caps when *η* is located in the intermediate value between 0 and 1, i.e., 0 *< η <* 1. The spherical caps for each *η* must have equal area as required by constraint constant area *A*_0_, implying 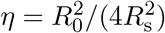.

**Figure 2.**
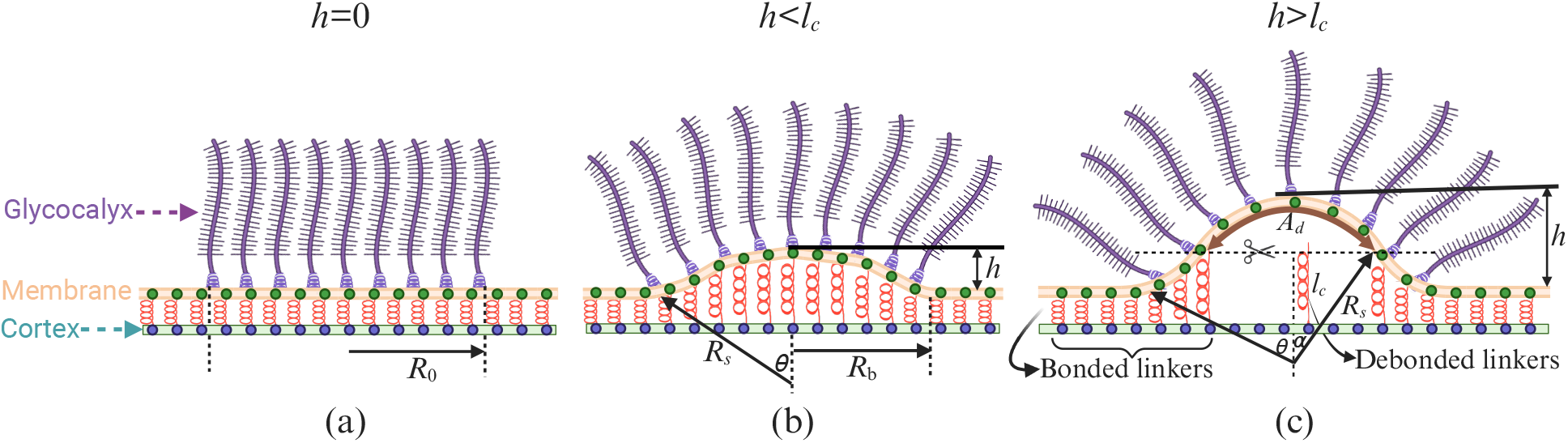
Schematic of the glycocalyx-membrane-cortex system for EV formation. (a) A patch of glycocalyx with inplane radius *R*_0_, represented as a layer of brush-like polymer, is anchored on an initially flat membrane. The cortex beneath the membrane is attached to the membrane through linker proteins, which are represented by springs. (b) The small deformation state where the membrane takes the shape of a spherical cap with its radius of curvature, base radius, opening angle, and height denoted by *R*_s_, *R*_b_, 2*θ*, and *h*. The linkers are attached to the membrane when the depth of the spherical cap is smaller than the maximum extension length of the membrane-cortex spring, i.e., *h < l*_c_. (c) In the case of *h > l*_c_, the spherical cap-shaped membrane is divided into two parts: the membrane-cortex adhesion region below the dashed line with bonded linkers, and the membrane-cortex detached region above the dashed line with debonded linkers. We denote the opening angle and area of the membrane-cortex detached region by 2*α* and *A*_d_. Panels are prepared using Biorender.com.

Given the attachment between the cell membrane and the actin cortex, membrane bending requires stretching and/or breaking of these adhesive connections. In addition to the glycocalyx and cortex, the line tension caused by the multiple phases on the membrane also affects membrane deformation. Thereby, the total free energy of the system is described as follows.

#### B. Energetics of the system

After phase separation, we assume that an initially flat circular domain, coated with glycocalyx, is embedded in a membrane matrix, its boundary experiences a line tension *λ* due to the unfavorable interactions at the interface between the glycocalyx-rich domain and the surrounding bare lipid layer. In this case, the domain edge energy that is proportional to the length of the perimeter and to the line tension is given by

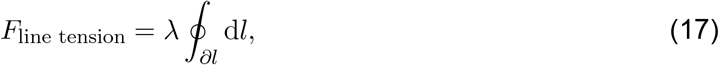

where *λ* denotes the strength of line tension along the domain boundary, and the integral is over the periphery line d*l* of the domain. We assume that the line tension is a constant. Consequently, by adding the line tension energy to Eq. (1) the total free energy of the glycocalyx-membrane-cortex system (*F*_tot_) becomes

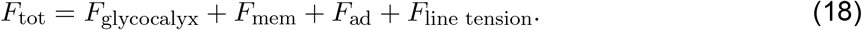

The first energy term on the right-hand side of Eq. (18) includes the elastic stretching of the polymer chains and the interactions between monomers in the polymer brush. Following Refs. (*53, 106*), this energy contribution originating from the glycocalyx polymers is given by

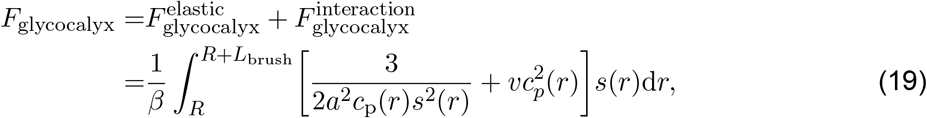

where *β* = 1*/*(*k*_*B*_*T*) and in which *k*_*B*_*T* is the thermal energy with *k*_*B*_ being the Boltzmann’s constant and *T* being the absolute temperature, *a* is the effective monomer size, *c*_p_(*r*) is the local monomer density profile along the thickness, *r* is the radial distance defined from the center of the spherical or cylindrical surface, *s*(*r*) is the area per chain at distance from the polymer grafting surface, and *v* is the second virial coefficient. Note that only pairwise monomer-monomer interactions with second virial coefficient *v* are considered. The detailed derivation of the individual energy component in Eq. (19) is provided in SI. 1. Using these expressions, we can write

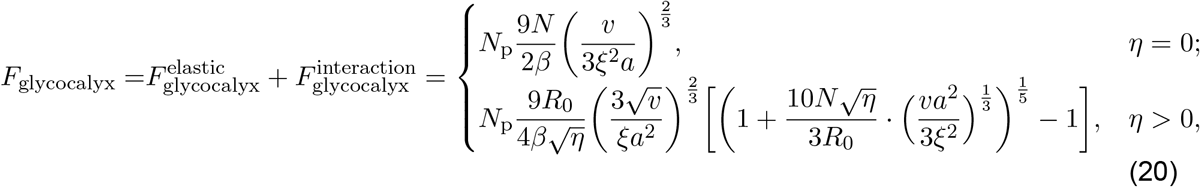

where *N*_p_ is the number of polymer chains grafted on the membrane, and *ξ* is the grafting distance of polymers on the cell membrane surface.

Here, in this model, the membrane geometry is restricted to spherical bud geometry since we are focused on EV formation, the two principal curvatures are given by *c*_1_ = *c*_2_ = 1*/R*_s_, the excess area is calculated as 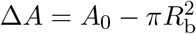, where 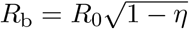 is the in-plane base radius, and the length of the domain boundary can be written as ∮*∂l* d*l* = 2*πR*_b_. Therefore, the elastic energy and the line tension of the membrane becomes

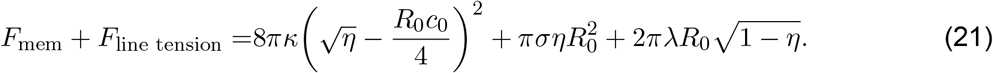

and the membrane-cortex adhesion energy appearing in Eq. (18) can be estimated as

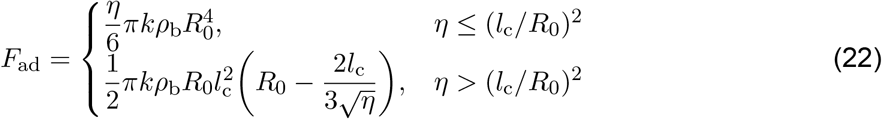

where *l*_c_ refers to the maximum length of the membrane-cortex adhesion bond. A complete derivation of this energy contribution can be found in SI. 2.

By substituting Eqs. (20), (21), and (22) into Eq. (18), we can obtain the total free energy of the system

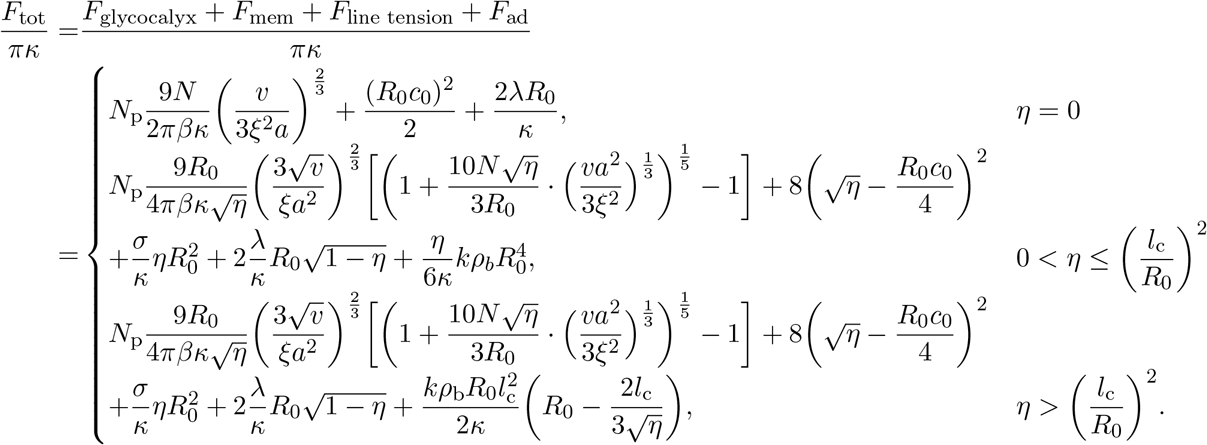

Here, we assume that our system is at equilibrium, implying that the system selects the membrane shape that minimizes the total free energy of the above equation.

#### III. Numerical implementation

To analyze the stability of the glycocalyx-membrane-cortex system, we numerically solve the eigen-value problem to obtain the eigenvalues which correspond to the linear growth rates of the perturbations. By displaying the corresponding dispersion relation functions Λ(*q*) at various degrees of control parameters, we can determine the stability behaviors.

For the equilibrium description model of the formation of EVs, we investigate the equilibrium geometry of the glycocalyx-membrane-cortex system under the constraint of fixed membrane patch area 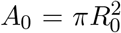 and the restriction of a spherical cap of membrane shape. We minimize the total free energy with respect to the shape parameter *η* to determine the stable (energetically most favorable) membrane shapes *η*_min_. Note that an analytic solution for *η*_min_ is not feasible due to the fact that the total free energy is nonlinear in the shape parameter *η*. Thus, we numerically determine *η*_min_ throughout this work by minimizing *F*_tot_. The values of parameters used in the model are summarized in Table 2 unless otherwise specified. The code is available on [https://github.com/Ranga-maniLabUCSD/Glycocalyx_Membrane_Cortex].

**Table 2.** Model parameter values.

| Parameter | Meaning | Estimated range | References | Default value |
| --- | --- | --- | --- | --- |
| $k$ | Spring constant | $\sim 10^{-4}$ N/m | (56, 57) | $2.5 \times 10^{-2} k_B T / \text{nm}^2$ |
| $\rho_0$ | Density of available linkers | $\sim 10^{14}-10^{16} \text{ m}^{-2}$ | (56, 57) | $10^{-4} \text{ nm}^{-2}$ |
| $k_{\text{on}}$ | Linker attachment rate | $\sim -10^3-10^4 \text{ s}^{-1}$ | (56, 57) | $10^4 \text{ s}^{-1}$ |
| $k_{\text{off}}^0$ | Free linker detachment rate | $\sim -0.1-10 \text{ s}^{-1}$ | (56, 57) | $10 \text{ s}^{-1}$ |
| $\delta$ | Linker bond length | $\sim 10^{-9} \text{ m}$ | (56, 57) | 1nm |
| $l_c$ | Critical length of membrane-cortex spring | $\sim 2-10 \text{ nm}$ | (56, 57) | 5 nm |
| $A_p$ | Polymer area | $\sim 100 \text{ nm}^2$ | (107) | $100 \text{ nm}^2$ |
| $D_{\text{diff}}$ | Polymer diffusion constant | $0.1-1 \text{ m}^2/\text{s}$ | (108) | $10^6 \text{ nm}^2/\text{s}$ |
| $M_h$ | Damping coefficient | $\sim 10^8 \text{ nm}^4/(k_B T \cdot \text{s})$ | (109) | $10^8 \text{ nm}^4/(k_B T \cdot \text{s})$ |
| $M_p$ | Mobility coefficient | $\sim 10^8 \text{ nm}^4/(k_B T \cdot \text{s})$ | (109) | $10^8 \text{ nm}^4/(k_B T \cdot \text{s})$ |
| $\phi_{p,\text{eq}}$ | Polymer area fraction | $\sim 0-1$ | (57) | 0.3 |
| $\omega$ | Polymer-membrane interaction | $\sim -10-10 k_B T$ | (74, 110) | $3 k_B T$ |
| $\chi_0$ | Polymer-polymer interaction | $\sim -10-10 k_B T$ | (109) | $-6 k_B T$ |
| $\gamma$ | Interfacial energy | $\sim 10^2 k_B T \cdot \text{nm}^2$ | (109, 110) | $200 k_B T \cdot \text{nm}^2$ |
| $\kappa$ | Bending rigidity | $10-400 k_B T$ | (111-114) | $10 k_B T$ |
| $\sigma$ | Membrane tension | $10^{-7}-10^{-3} \text{ N/m}$ | (112, 115, 116) | $0.012 k_B T / \text{nm}^2$ |
| $\lambda$ | Line tension | $0-100 \text{ pN}$ | (112, 117, 118) | $0.5 k_B T / \text{nm}$ |
| $R_0$ | Patch radius | $20-100 \text{ nm}$ | (93, 112) | 100 nm |
| $\xi$ | Grafting distance | $10-100 \text{ nm}$ | (53, 119) | 15 nm |
| $a$ | Effective monomer size in the glycocalyx | $10-20 \text{ nm}$ | (50, 53, 93, 120-122) | 10 nm |
| $N$ | Number of effective monomers in a brush | $10-50$ | (50, 53, 93, 122) | 20 |
| $c_0$ | Spontaneous curvature | $0-0.1 \text{ nm}^{-1}$ | (53) | $0.02 \text{ nm}^{-1}$ |

## Results and Discussions

### Different stability behaviors

In this section, we focus on stability analysis. In order to classify the types of instability exhbited by the system, we begin by examining the dispersion relations Λ(*q*), i.e., growth rates as a function of the wave number at different values of control parameters. We plot the dispersion relations Λ(*q*) at different values of *ρ*_b_ in Fig. 3(a). For low linker density (*ρ*_b_ = 0.01), there exists an instability band that starts at *q*_*−*_ and lasts until *q*_+_, with the most unstable mode at *q*_*−*_ *< q*_max_ *< q*_+_. As we decrease the linker density, the instability band narrows to only a most unstable mode *q*_c_ = *q*_max_ when the linker density reaches a threshold value 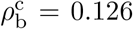. Here, *q*_c_ corresponds to the critical wavenumber where at the onset of instability a maximum of Λ(*q*) touches zero, *q*_*±*_ represents the marginal wavenumbers where Re(Λ_max_) crosses zero above the onset, and *q*_max_ denotes the fastest growing wavenumber at where Re(Λ_max_) has a maximum above the onset. A density beyond 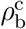, i.e., 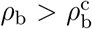, leads to the disappearance of the instability band, which indicates that the system is stable in any mode and the initial perturbations decay exponentially. This type of dynamic behavior corresponds to the conserved Turing-type (cT-type) instability (*105, 109*) which occurs for some range of wavenumbers (Re(Λ_*i*_) *>* 0), *q*_*−*_ *< q < q*_+_, and where there are non-oscillatory unstable modes (Im(Λ_*i*_) = 0), while the zero wavenumber remains stable. In this region, the amplitudes of these modes grow exponentially from small initial perturbations but do not oscillate or propagate on the membrane surface. The physical mechanism behind this type of instability is that, when the linker density is low, the perturbed homogeneous glycopolymer starts to aggregate and undergoes phase separation and protrusions start to appear on the initially flat membrane. This indicates the relevant biological prediction that decreases in membrane-cortex detachment can give rise to an unstable state and initiate cell protrusions. Furthermore, we present the band of unstable wavenumbers (*q*_*−*_, *q*_+_) as a function of the linker density *ρ*_b_, as shown in Fig. 3(b). For 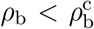, the dispersion relation exhibits a cT-type instability, which exists in some ranges of modes, *q*_*−*_ *< q < q*_+_. Above a threshold value 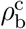, the band of unstable modes disappears and the system remains steady state.

**Figure 3.**
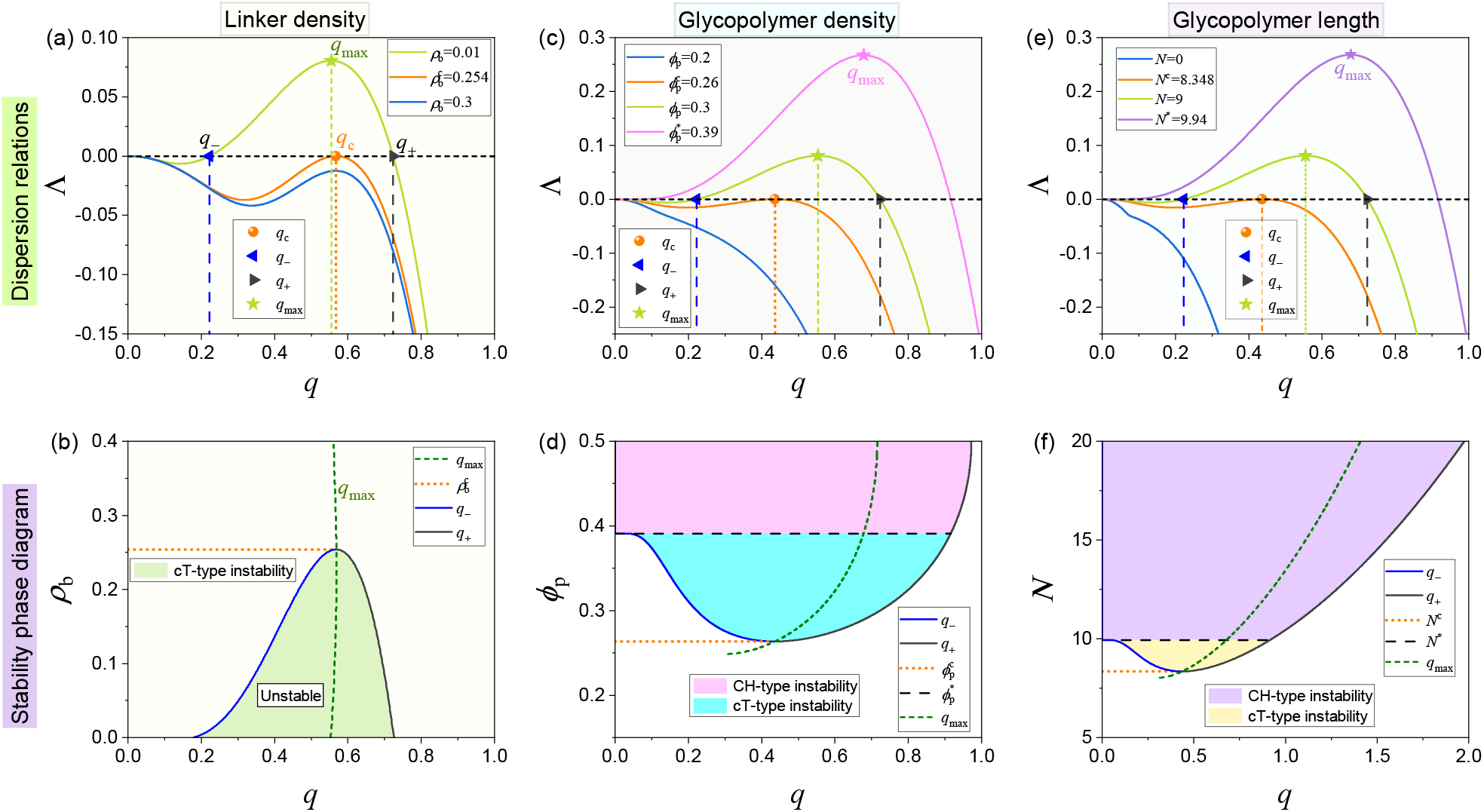
Linear stability analysis of the system. (a) Dispersion relation functions Λ_*q*_ at different values of *ρ*_b_. (b) Band of unstable modes (*q*_*−*_, *q*_+_) in the plane spanned by wavenumber *q* and control parameter linker density, *ρ*_b_. (c) Dispersion relations for different values of *ϕ* _p_. (d) Band of unstable modes (*q*_*−*_, *q*_+_) in the *q − ϕ* _p_ plane. (e) Growth rate for different values of the control parameter *N*. (f) Stability diagram of the *q− N* plane illustrating the band of unstable wavenumbers in its dependence on the length of glycopolymer *N*.

In addition to the effect of membrane-cortex attachment, the properties of the glycocalyx also affect the stability behaviors of the system. Figure 3(c) shows the growth rate as a function of wavenumber at different values of *φ*_p_. At low glycocalyx density (see blue curve), the largest growth rates Λ(*q*) for all modes are negative except the neutral mode at zero wavenumber, i.e., Λ = 0 at *q* = 0. Upon increasing the glycocalyx density to a critical value 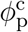 (see orange curve), we observe that a nonzero marginal wavenumber appears where the maximum of Λ(*q*) touches zero, this implies the onset of instability. Then, above this critical value 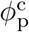 results in an instability band which starts at *q*_*−*_ and last until *q*_+_ (see light green curve). This instability pattern corresponds to the conserved Turing-type (cT-type) instability. While as the glycocalyx density above another threshold value 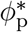, the dispersion curve exhibits an instability band starting at *q* = 0 and ending at some *q* = *q*_+_, with the most unstable mode at 0 *< q*_max_ *< q*_+_ (see magenta curve). This type of instability is referred to as Cahn-Hilliard-type (CH-type) instability (*109, 123*), which is a long-wavelength instability with the band extending to the zero wave vector (*q*_*−*_ = 0). Finally, we present the stability diagram of the *q−φ*_p_ plane to demonstrate the band of unstable wavenumbers (*q*_*−*_, *q*_+_) as a function of the glycocalyx density *φ*_p_, as shown in Fig. 3(d). For 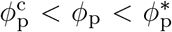, the system displays a cT-type instability, while when 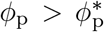, the dispersion relation shifts from a cT-type instability to a CH-type long-wavelength instability. This means that when the glycocalyx density is high enough, the glycocalyx can undergo phase separation on the membrane surface and initiate EV formation.

In addition to grafting density, length of the glycopolymers is another important feature of the glycocalyx layer on the cell membrane surface. To probe the influence of glycopolymer length on the stability behaviors of the system, the analysis of dispersion relations for various numbers of monomers *N* is presented in Fig 3(e). For short glycopolymer lengths (i.e., *N < N* ^c^), the system does not demonstrate an instability. Similarly, depending on the length of glycopolymer, we observe two types of instabilities. For glycopolymer lengths in the regime *N* ^c^ *< N < N* ^*\**^, the instability behavior shows a cT-type instability, which exists for some range of modes, *q*_*−*_ *< q < q*_+_. However, further increasing the length of glycopolymer to a threshold value *N* ^*\**^ leads to a extending of the instability band (*q*_*−*_, *q*_+_) which corresponds to a reduction of the left marginal wavenumber *q*_*−*_ to zero. As a result, above *N* ^*\**^, a change in the type of instability from cT to CH type occurs, where the dispersion relation exhibits an instability band starting at *q* = 0 and ending at some *q* = *q*_+_, with the most unstable mode at 0 *< q*_max_ *< q*_+_. Therefore, varying the length of glycopolymer provides a practical strategy to tune the aggregation and phase separation on the membrane and to regulate membrane curvature generation. Similarly, we constructed a two-dimensional phase diagram on the *q − N* plane to characterize the band of unstable modes in its dependence on the length of glycopolymer *N*, as demonstrated in Fig. 3(f). Our results show that the dispersion relation shifts from a cT-type instability to a CH-type long-wavelength instability as *N* reaches a critical value *N* ^*\**^. This suggests that long glycopolymers have the tendency to destabilize the system and trigger membrane-cortex release and facilitate the onset of EV formation.

The stability analysis reveals the regimes in the control parameter space where the system is stable and where it is unstable. The unstable regimes exhibit two types of instabilities: cT-type instability and CH-type instability, and there the glycopolymer undergoes aggregation and the membrane protrusions appear. Analogous to Fig. 3, the influence of spontaneous curvature and membrane properties (i.e., membrane tension) on the stability of the system is presented in Fig. S2 (see SI. 3). Next, we will employ the equilibrium description model to probe the conditions under which the EV can be successfully formed.

### Effects of glycocalyx and membrane-cortex adhesion on EV formation

In this section, we numerically calculate the total energy for different shape parameters *η* ranging from 0 to 1 based on the equilibrium description model of the formation of EVs. The global energy minimum corresponds to the optimal *η*_min_ that gives the equilibrium geometry of the cell membrane patch. Here, we first investigate the influence of the level of membrane-cortex protein linkers on energy as well as membrane shape. To quantify the effect of membrane-to-cortex attachment on the initiation of EV formation, we focus on the total energy profile as a function of the shape parameter *η* for a series of fixed linker density *ρ*_b_, as shown in Fig. 4(a). Immediately, we observe that there exists a global energy minimum (solid dot) which corresponds to the formation of a spherical EV (0 *< η <* 1 or a partial blebbing state). As *ρ*_b_ increases (from *ρ*_b_ = 10^*−*4^ nm^*−*2^ to *ρ*_b_ = 3 *×* 10^*−*2^ nm^*−*2^), the global stable state slightly shifts to a shape corresponding to a smaller *η*. Interestingly, we observe that the stable partial blebbing state shifts to a stable no blebbing state with increasing *ρ*_b_ to a high level (refer to the curve with *ρ*_b_ = 4 *×* 10^*−*4^ nm^*−*2^), indicating that the membrane remains attached to the cortex and undeformed when the density of cortex-membrane protein linkers is higher enough. Note that the initial flat membrane, in order to grow up to a spherical EV, needs to overcome an energy barrier Δ*F*.

**Figure 4.**
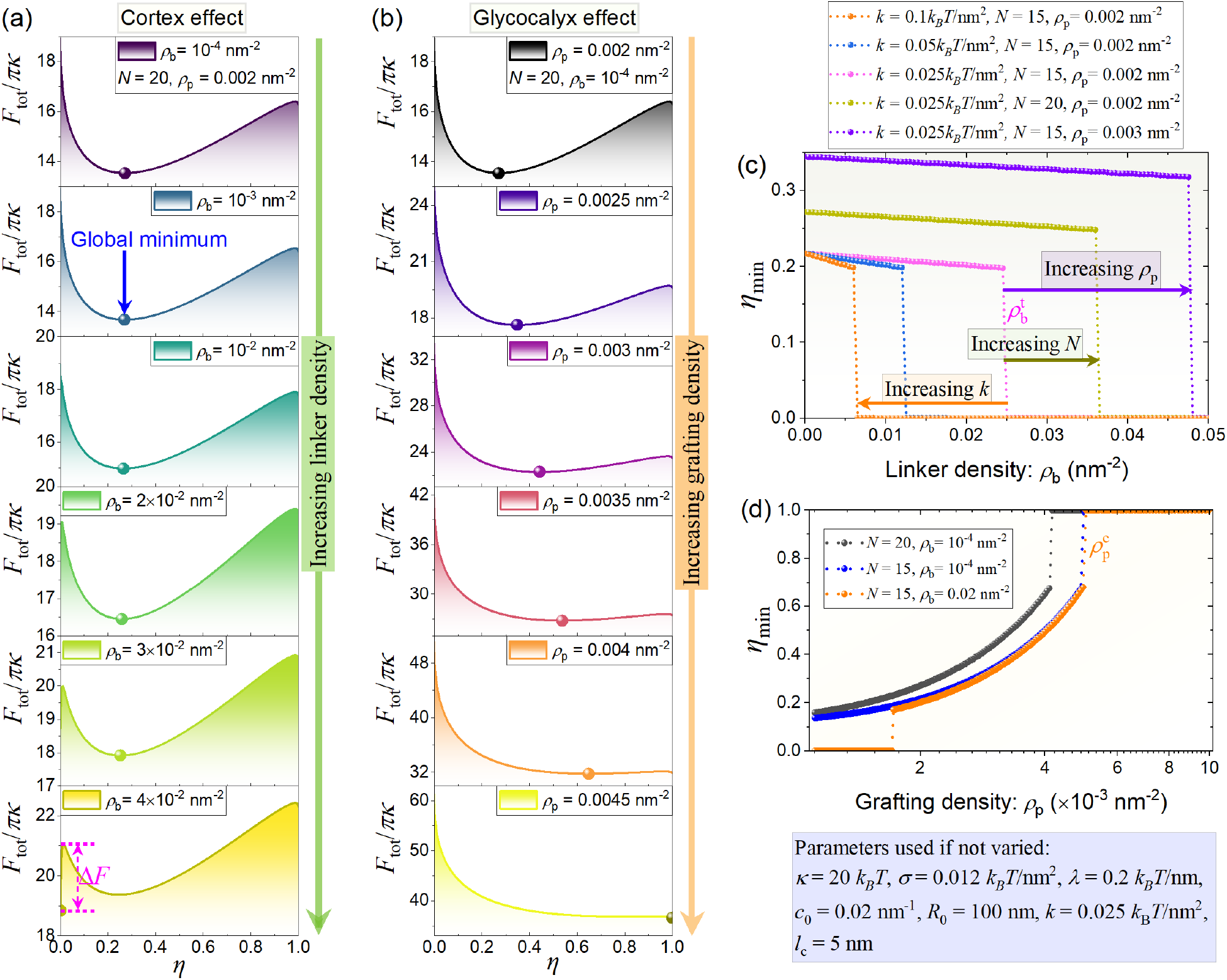
Total energy profile as a function of shape parameter *η* for different values of (a) linker density *ρ*_b_ and (b) glycopolymer density *ρ*_p_. The dependence of minimum shape parameter *η*_min_ on (c) linker density *ρ*_b_ and (d) glycopolymer grafting density *ρ*_p_.

Apart from membrane-cortex adhesion, the properties of glycocalyx such as grafting density and length also has a big impact on EV formation. In Fig. 4(b), we present the total energy profile as a function of the shape parameter *η* for a sequence of fixed glycopolymer grafting density *ρ*_p_. Similar to Fig. 4(a), a global energy minimum also exists (see the solid dot) and the stable state still corresponds to the partial blebbing state (*η <* 1) at low polymer grafting density (i.e., the curve with *ρ*_p_ = 0.002 nm^*−*2^). As the grafting density increases, the stable partial blebbing state shifts to a stable state with a full EV formed (refer to the curve with *ρ*_p_ = 0.0045 nm^*−*2^). This implies that a higher grafting density of glycopolymer on the cell membrane is necessary to achieve a full EV.

Next, to intuitively visualize the effect of membrane-cortex attachment, we focus on the stable state corresponding to the optimal shape parameter *η*_min_. Figure. 4(c) shows the dependence of *η*_min_ on linker density at different linker stiffness, polymer length, and grafting density. We note three main effects of varying linker density. First, all curves in Fig. 4(c) illustrate that the optimal shape parameter *η*_min_ shows a sharp jump (0 *→ η*_min_ *∈*(0, 1)) at a critical onset value, *ρ*^t^, a characteristic of discontinuous transition. Such a transition indicates that reducing the linker density to a specific threshold is necessary to trigger the detachment of the membrane from the cortex. In other words, the membrane remains flat (*η* = 0) and the shape transformation is unsuccessful when the linker density is above a certain value 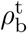, which reflects that increasing the level of membrane-cortex protein linkers and enhancing the membrane-cortex attachment reduces the formation of EV. This implies that local depletion or disruption of membrane-cortex protein linkers is needed for membrane deformation initiation to occur. This prediction is consistent with experimental evidence from previous studies (*64*), where Welf et al. (*64*) have shown that plasma membrane detachment from actin is needed to initiate cell protrusion and increased actin-membrane attachment reduces protrusion. Second, this critical onset value depends on the linker stiffness (see the orange, blue and magenta curves), glycopolymer grafting density (comparing the dark yellow curve with the magenta curve) and polymer length (see the dark yellow curve and the violet curve). The curves in Fig. 4(c) show that the decreasing of linker stiffness or increasing glycopolymer grafting density and length results in a shift of the threshold 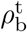, which triggers the discontinuous transition from *η* = 0 to *η <* 1. Third, at low stiffness, the optimal shape parameter *η*_min_ exhibits a gradual reduction as we increase the linker density *ρ*_b_. In contrast, the variation of the optimal shape parameter *η*_min_ becomes more pronounced and is accompanied by a steeper slope with the increasing of linker stiffness *k*. This effect can be intuitively understood by considering the role of the membrane-cortex adhesion energy in total energy. Since this adhesion energy opposes membrane bending, a higher energy penalty is incurred when membrane deformation occurs. As a result, it becomes more difficult to detach the cell membrane if the cortex contains a higher linker density with stronger stiffness. We also observe that increasing the glycopolymer grafting density or length raises the optimal shape parameter, shifting curves *η*_min_ upward with increasing *ρ*_p_ or *N* (comparing the magenta curve with the dark yellow curve or the violet curve). This rise in the optimal shape parameter indicates that the glycocalyx serves as a driving mechanism for the formation of EVs.

Finally, we plot the optimal shape parameter *η*_min_ against the glycopolymer grafting density *ρ*_p_ at different lengths of the polymer chain (see the black and blue dotted curve) and linker density (see the blue and orange dotted curve), as shown in Fig. 4(d). We found that the optimal shape parameter *η*_min_ exhibits a snap-through jump at a threshold 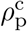 at high grafting density, a characteristic of first-order transition. This suggests that the membrane geometry transits from a partially spherical EV to a full sphere EV. Comparison of the black and blue curves reveals that shortening the polymer chain length lowers the threshold grafting density required to trigger the formation of a fully spherical membrane geometry. Moreover, comparing the blue curve and the orange curve, we observe that, at low surface coverage of glycopolymers, (i) the undeformed state occurs when the liker density is higher enough; (ii) the optimal shape parameter *η*_min_ undergoes a sharp jump (also indicates a first-order transition) as the grafting density reaches a critical value. In other words, a critical value of grafting density, 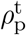 is needed to initiate membrane detachment from an initially flat membrane. However, the blue and orange curves in Fig. 4(d) almost overlap, indicating that increased linker density has minimal effect on *η*_min_ and 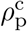 once membrane detachment is onset.

Therefore, we find that there exist two threshold grafting densities, one is the critical value required to initiate membrane detachment from a flat membrane, and the other one corresponds to the critical value that triggers a transition from a spherical EV to a full sphere EV.

### Phase diagrams for glycocalyx-driven EV formation

To characterize the interrelated effects of glycopolymer grafting density and length on the formation of EV, we constructed two-dimensional phase diagrams on the (*ρ*_p_–*N*) plane in four different cases, as shown in Fig. 5. For both low and high membrane tension, each situation corresponds to two distinct cases: (i) low linker density (see Fig. 5(a) and (c)); and (ii) high linker density (see Fig. 5(b) and (d)). Figure 5 illustrates three regimes corresponding to undeformed membrane (regime I), spherical cap (regime II), and full sphere (regime III), with the color representing the value of the optimal shape parameter *η*_min_. The two-dimensional phase diagrams in the (*ρ*_p_–*N*) plane show a common feature that as the glycocalyx density or polymer chain length increases, the shape of the cell membrane undergoes a transition from a spherical cap to a full sphere when the values of *ρ*_p_ and *N* exceed certain thresholds. A comparison between Fig. 5(a) and Fig. 5(c) reveals that the glycocalyx can drive the membrane to deform into a full EV (or a complete sphere morphology) under the condition of crossing the threshold of the grafting density and polymer length. Whereas high membrane tension leads to a narrow regime III and shifts the boundary curve to the right (see the green curve), which indicates that a higher threshold of the grafting density and polymer length is required to regulate the formation of full sphere EV. When comparing Fig. 5(a) (or (c)) and Fig. 5(b) (or (d)), we observe that an increase in the level of linker density will result in the emergence of the non-deformed regime (flat membrane regime). This validates the prediction presented in Fig. 4, namely, that membrane detachment cannot occur if the glycocalyx grafting density is below a certain threshold. Furthermore, the comparison of regime I between Fig. 5(b) and Fig. 5(d) illustrates that the non-deformed regime becomes broader, and the boundary curve (black curve) separating regime I from regime II shifts to the right in the case of high membrane tension. Figure 5(e) shows several typical cell membrane morphologies identified in the phase diagrams. We compare the computed EV geometries with those observed in Refs. (*124, 125*) for distinct types of cancer cells (Fig. 5(f)). The right column in Fig. 5(f) shows that aggressive tumor cells secrete spherical microvesicles (MVs are indicated by arrows) at their plasma membrane for long-range delivery of cargoes (*124*). The left column in Fig. 5(f) demonstrates that, to mediate intercellular communication, the production of MVs (MVs are denoted by arrowheads) is enhanced when breast cancer cells are exposed to decreased oxygen availability (hypoxia) (*125*). The EV shapes observed in the experiments bear resemblance to spherical geometry, which supports the applicability of our model in predicting the formation of EVs.

**Figure 5.**
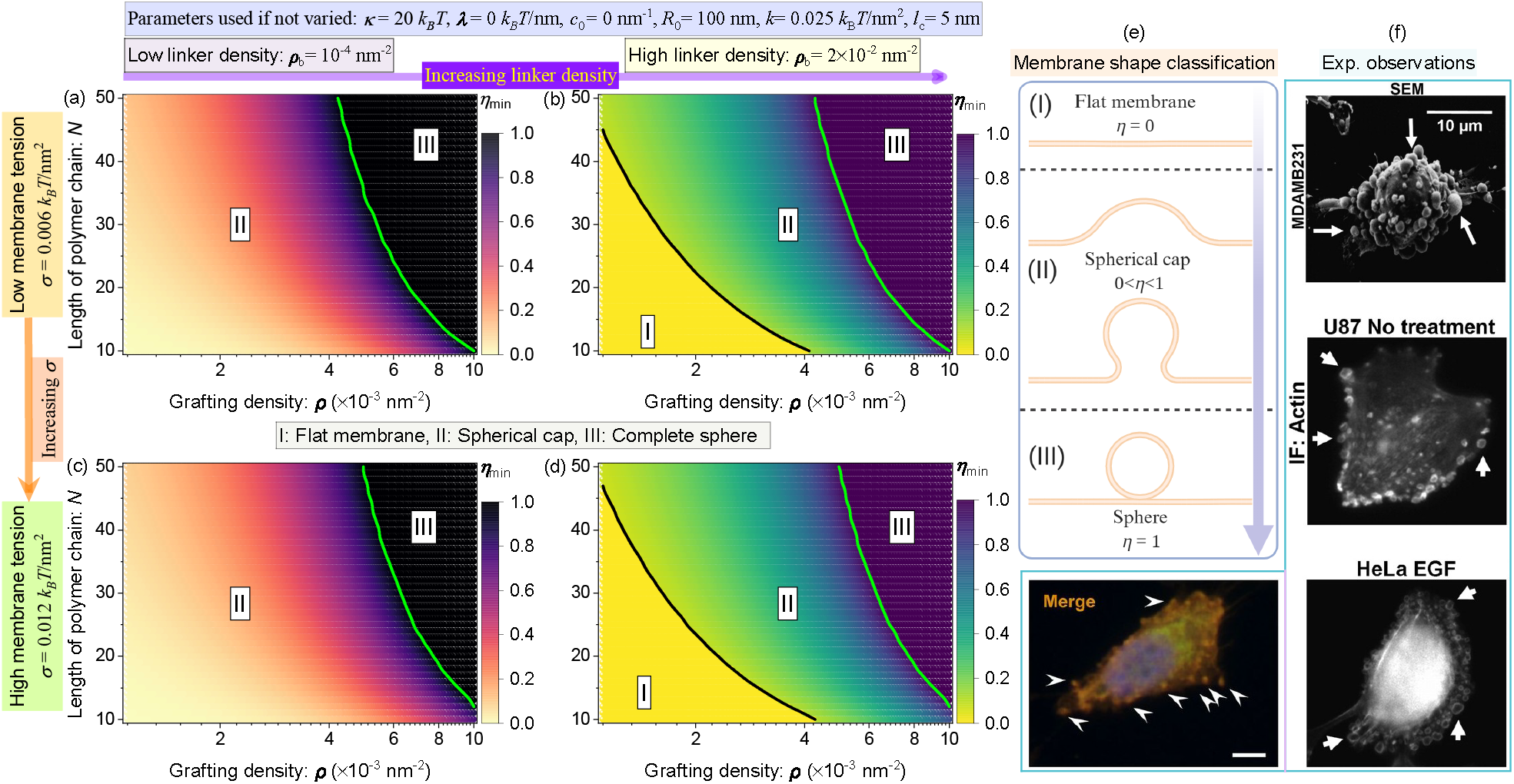
Heatmaps of optimal shape parameter *η*_min_ as a function of glycocalyx grafting density, *ρ*_p_, and polymer chain length, *N*, are presented for various scenarios. Panels (a) and (b) correspond to low membrane tension, while panels and (d) correspond to high membrane tension. Specifically, (a) and (c) represent low linker density, and (b) and represent high linker density. (e) Sketch of various membrane morphologies correspond to flat, spherical cap, and sphere shapes. (f) Observed microvesicles (MV), a subtype of EVs, releasing from different types of cancer cells (M. A. Antonyak et al., 2011 and T. Wang et al., 2014) (*124, 125*). (f) Reprinted from Refs. (*124, 125*); permission pending.

In summary, membrane-cortex adhesion and glycocalyx properties, such as its density and length, play a crucial role in determining the equilibrium geometry of the cell membrane. The boundary curves that separate the distinct states shift when linker density is altered. Also, to explore the influence of spontaneous curvature and line tension on EV formation, a two-dimensional phase diagram on the (*λ − c*_0_) plane is constructed to characterize the interrelated effects of line tension and spontaneous curvature on the shape of the membrane, as shown in Fig. S3 (see SI. 4).

Our results demonstrate that EV formation is more easily achieved in the presence of line tension and spontaneous curvature.

### Influence of membrane-Cortex adhesion and glycocalyx coverage on EV formation

It has been reported that EVs exhibit a broad size distribution, typically ranging from 30 to 1000 nanometers (*22, 23*). Therefore, it is necessary to understand the role of the glycocalyx coating area in determining EV formation. We first perform the analysis of total free energy for different values of patch radius *R*_0_. In Fig. 6(a), we plot the total free energy landscape for different *R*_0_. A direct comparison of the global energy minimums for *R*_0_ = 50 nm, *R*_0_ = 100 nm, and *R*_0_ = 300 nm reveals that a larger glycocalyx grafting area tends to induce the formation of EV. However, further increase of *R*_0_ leads to the formation of EV more unfavorable (see the curves for *R*_0_ = 500 nm, *R*_0_ = 700 nm, and *R*_0_ = 900 nm, respectively). This implies that the effects of the glycocalyx grafting area on EV formation show a non-monotonic behavior. Figures. 6(b)-(d) show the dependence of *η*_min_ on *R*_0_ for a series of fixed linker density, glycocalyx grafting density, and glycopolymer length. At low linker density, the blue curve in Fig. 6(b) shows that *η*_min_ depends on *R*_0_ in a non-monotonic manner which increases first and is followed by a monotonic drop-off. However, the minimum shape parameter *η*_min_ undergoes a discontinuous reduction to zero at a critical *R*_0_ (see open star) under the circumstance of high liker density. Here, we define the optimal radius 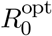 (solid star) at which *η*_min_ reaches maximum.

**Figure 6.**
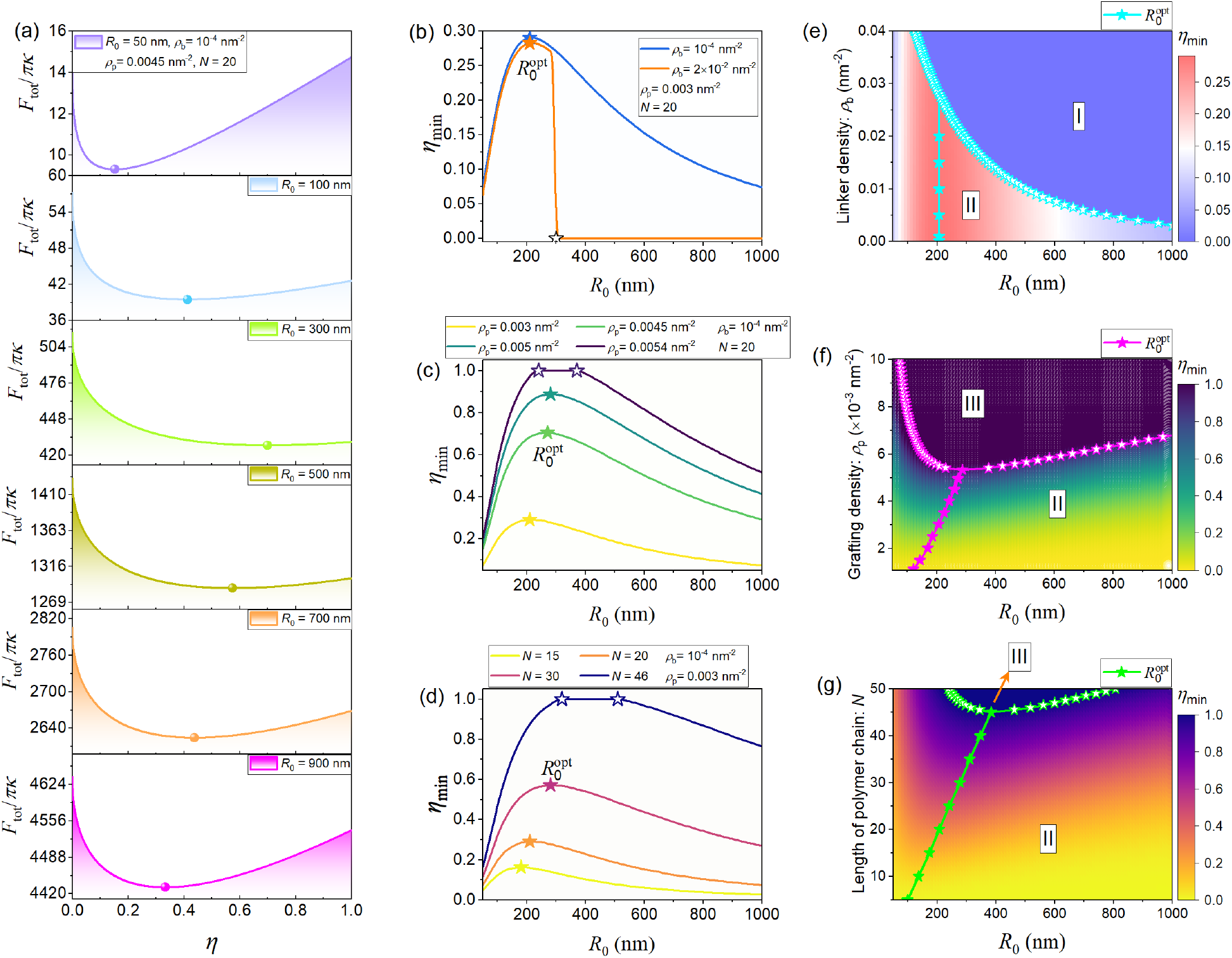
(a) Total free energy profile as a function of shape parameter *η* for different values of *R*_0_. The dependence of the optimal shape parameter *η*_min_ on *R*_0_ at different (b) linker density *ρ*_b_, (c) glycocalyx grafting density *ρ*_p_, and (d) glycopolymer length *N*. Phase diagrams present the optimal shape parameter *η*_min_ as a function of *R*_0_ and (e) linker density *ρ*_b_, (f) glycocalyx grafting density *ρ*_p_, and (g) glycopolymer length *N*.

By inspecting the dependence of the optimal shape parameter *η*_min_ on *R*_0_ for a series of fixed glycocalyx grafting density *ρ*_p_ (see Fig. 6(c)) and glycopolymer length *N* (see Fig. 6(d)), we observe a analogous variation trend which is *η*_min_ showing a monotonous rise as 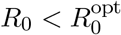 while a gradual reduction for 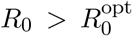. Notably, when the glycocalyx grafting density or glycopolymer length reaches a critical value, the variation of the optimal shape parameter *η*_min_ is accompanied by a plateau in some range of radii (see the band between the open stars in Figs. 6(c) and (d)), indicating the successful formation of a sphere EV.

Furthermore, we scan the phase spaces of *R*_0_ *− ρ*_b_, *R*_0_ *− ρ*_p_, and *R*_0_ *− N* in numerical calculations to determine their influence on the optimal shape parameter *η*_min_, as shown in Figs. 6(e)-(g), where the magnitude of *η*_min_ is shown as a color map. Note that all features of Fig. 6(b), (c), and (d) can be captured by Fig. 6(e), (f), and (g), respectively. Additionally, Fig. 6(e) shows that two regimes corresponding to non-deformed (regime I: *η* = 0) and inflated (regime II: 0 *< η <* 1) states are identified. While Figs. 6(f) and (g) include regime II and regime III (*η* = 1) due to that the glycocalyx serves as a driving mechanism for EV formation. The open star dotted line that separates regime I from regime II represents the critical values. Below this critical line implies that membrane detachment is initiated and an inverse dependence relationship is displayed between the critical *R*_0_ and *ρ*_b_. Intriguingly, the optimal radius slightly depends on linker density (see the solid star dotted line). Remarkably, in contrast to Fig. 6(e), the dependence of 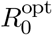 on glycocalyx grafting density and glycopolymer length in Fig. 6(f) and Fig. 6(g) shows an almost linear proportional relationship (see the solid star dotted line). While the critical curves that separate regime II from regime III exhibit a non-monotonic behavior. Two more phase diagrams on the (*R*_0_ *−ρ*_b_) plane at different glycocalyx grafting density and glycopolymer length are presented in Fig. S4 (see SI. 5) to show that increasing the grafting density and polymer length can narrow regime I as compared to Fig. 6(e).

In summary, our results demonstrate that membrane-cortex adhesion inhibits membrane detachment, while glycocalyx plays a positive role in assisting EV formation. Inspecting the role of the glycocalyx grafting area, we can infer that the formation of EVs becomes more favorable as *R*_0_ reaches the optimal value 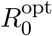.

### Size-dependent probability of EV formation

Furthermore, to explore the possibility of generating a spherical EV, indicating that the EV is in a state with *η* = 1, we assume that the morphology of the EV is governed by a Boltzmann distribution. As a result, the probability of forming a closed EV can be obtained from statistical mechanics, which is given by

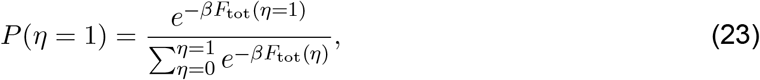

where *F*_tot_ is given by Eq. (18). We numerically evaluate *P* (*η* = 1) in Eq. (23) to predict the probability of generating a spherical EV. Figures 7(a) and 7(b) show the *P* (*η* = 1) as a function of *R*_0_ for different values of the glycocalyx grafting density and polymer length. We observe that, for glycocalyx with a sparse grafting density and a short length (i.e., *ρ*_p_ = 0.005 nm^*−*2^ in 7(a) and *N* = 15 nm^*−*2^ in 7(b)), the possibility of an EV in the closed state is nearly zero, which implies that generating a closed EV is almost impossible. Compared to the flat curve, the rest of the curves in Figs. 7(a) and 7(b) demonstrate that the larger the values of grafting density and polymer length, the greater the likelihood of generating a closed EV. In addition, we also note that the width of the spectrum of 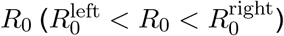 spanned by the nonzero probability distribution profile becomes wider with increasing glycocalyx grafting density or polymer length, indicating that a wide range of EV sizes can be produced. Based on the probability distribution profile, we also defined a most possible radius 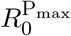 (solid dot) at which the odds of generating a closed EV has the highest value *P*_max_.

**Figure 7.**
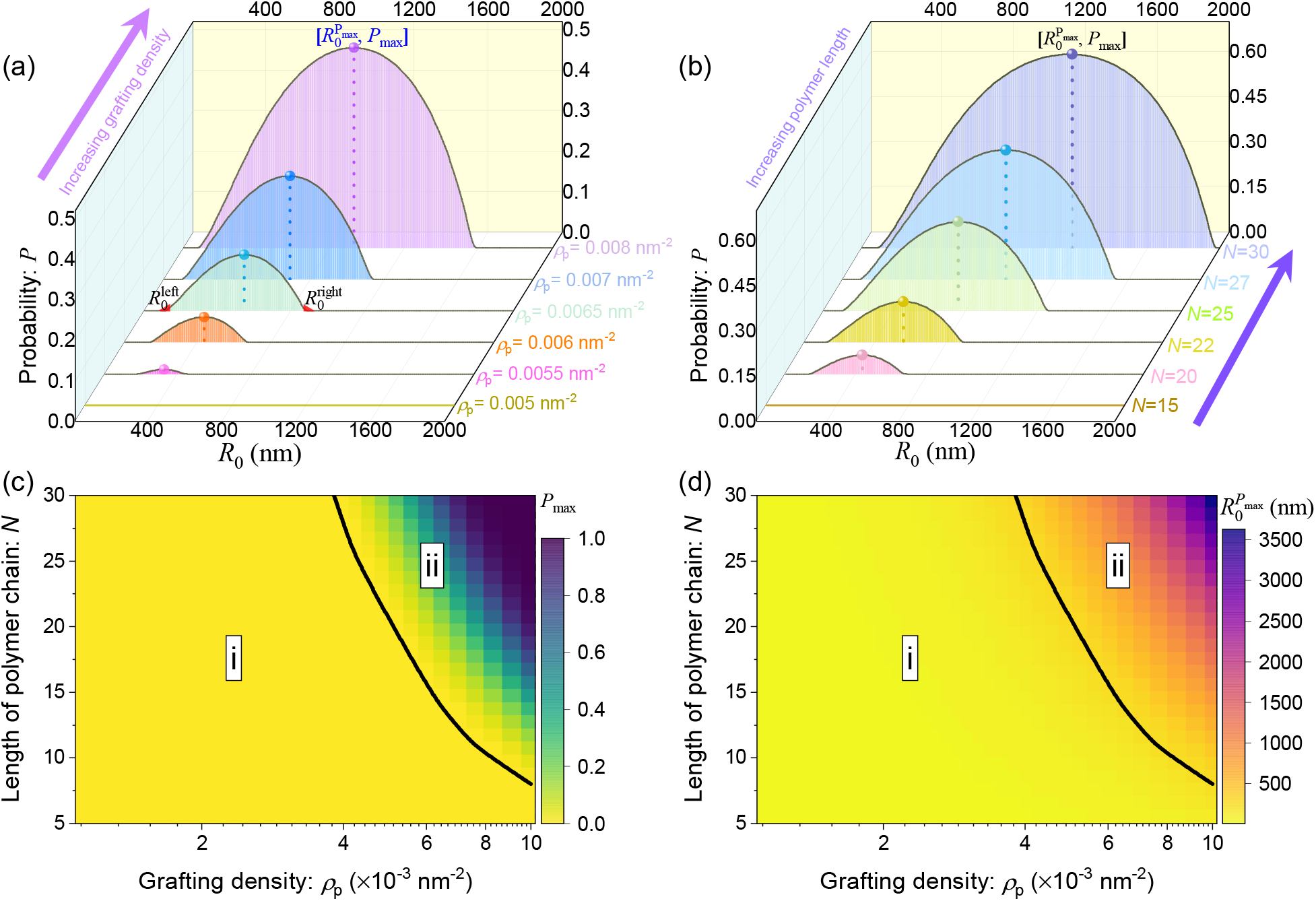
Probability of generating a sphere EV as a function of *R*_0_ for a set of fixed values of (a) glycocalyx grafting density *ρ*_p_, and (b) glycopolymer length *N*. Heatmap of (c) maximum probability of generating a closed EV, *P*_max_, and (d) corresponding 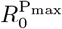 as a function of glycocalyx grafting density *ρ*_p_, and glycopolymer length *N*.

The interrelated effects of glycocalyx grafting density and polymer length on 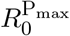 and *P*_max_ are summarized in Figs. 7(c) and 7(d). Specifically, depending on the properties of the glycocalyx, two regimes corresponding to *P*_max_ = 0 and *P*_max_ ≠ 0 cases are identified in Figs. 7(c), in which the color indicates the maximum probability of forming a closed EV. In regime i, it suggests that forming a sphere EV is nearly impossible. In contrast, in regime ii, the feasibility of producing a closed EV is more higher at denser and longer glycocalyx. The value of 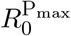 corresponding to regime ii is presented in Fig. 7(d). Similarly, Fig. 7(d) illustrates that larger EV can be produced at higher glycocalyx grafting density and longer polymer length. Therefore, our results predict that within the range of *ρ*_p_ and *N* relevant for EV, different combination of *ρ*_p_ and *N* tend to yield distinct EV sizes. This suggests that cells may self-regulate glycocalyx properties to produce a heterogeneous collection of EV size. Our prediction can be supported by previous experimental observations in Ref. (*53*) (refer to Fig. 6) where Shurer at al. observed that massive concentrations of particles ranging in size from approximately 100 nm to 400 nm can be secreted from Muc1-42TR-expressing cells. Importantly, they verified that the released particles are membrane vesicles with glycocalyx grafted on their surface by using cryo-transmission electron microscopy (cryo-TEM) analysis, and the removal of the glycocalyx significantly reduces the production of vesicles. Our results in Fig. 6 and Fig. 7 also imply that reducing glycocalyx grafting density will result in unsuccessful EV formation.

### Critical values required for initiating membrane detachment and EV formation

In Figs. 4, 5, and 6, we have identified two distinct critical conditions: one for the onset of glycocalyx-driven membrane detachment, and another for the regulation of the formation of full EV states. Based on our theoretical model, we expect that several parameters can affect these two critical conditions. To gain more insight into the effects of linker density on the critical condition that initiates membrane detachment, we plot the dependence of critical value of different parameters to initiate membrane detachment on linker density in Fig. 8(a). Intriguingly, we find that 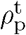 and *ρ*_b_ exhibit a logarithmic linear dependence, and *N*_t_, *R*_0,t_, *λ*_t_, *c*_0,t_, and *σ*_t_ are almost proportional to *ρ*_b_ linearly. Then, focusing on the dependence of the critical grafting density on various parameters in Fig. 8(b), we also find that 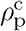 depends on *λ, c*_0_, and *σ* in a nearly linear manner, while 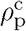 and *N* display a logarithmic linear dependence. Notably, the variation of the linker density rarely affects 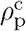 and the dependent relation between 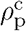 and *R*_0_ exhibits a non-monotonic behavior. In general, these observations predict that an increase in the level of linker proteins requires a higher grafting density (or a longer polymer length, a larger line tension, spontaneous curvature, a smaller grafting area, and a lower membrane tension) to initiate membrane delamination from the cortex. Longer polymer chains and looser membranes, along with the presence of line tension and spontaneous curvature, reduce the critical grafting density of the polymer required to trigger the formation of a spherical membrane structure.

**Figure 8.**
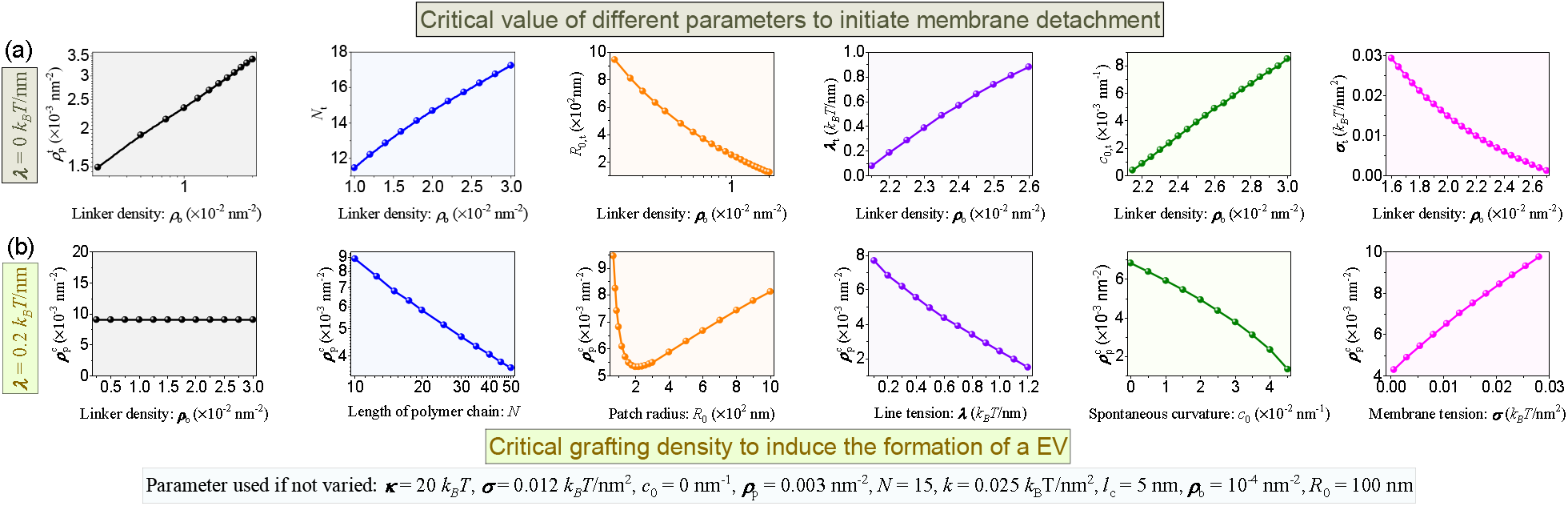
The dependence of (a) critical value of different parameters to initiate membrane detachment on linker density; (b) critical grafting density on different parameters, including linker density, length of polymer chain, patch radius, line tension, spontaneous curvature, and membrane tension.

## Discussion and Conclusion

Despite the fact that many experimental studies have shown that glycocalyx can regulate membrane morphologies (*53*) and that local depletion of actin-membrane bonds is needed for the initiation of membrane protrusions (*64*), we know surprisingly little about the critical conditions for the onset of membrane detachment and the formation of EVs. From a modeling perspective, we developed a comprehensive biophysical model that incorporates various factors to provide deeper insight into how the interplay between membrane-cortex adhesion and glycocalyx regulates EV formation. Based on our theoretical results, we make the following experimentally relevant predictions. The reduction of membrane-cortex linkers is necessary for the onset of membrane bending, which is qualitatively consistent with the experimental results observed by Welf et al. (*64*). This could be applied to understand the correlation between the level of membrane-cortex linker molecules and the potential for EV or bleb formation. For example, reduced blebbing has been experimentally observed in zebrafish germ cells (*126*), A375 melanoma cells (*127*), and mast cells (*128*) as the level or activity of ERMs increases. Bleb formation at the leading edge of the cell is facilitated due to the reduction of membrane-to-cortex attachment at the cell front, while the level of the actin-membrane linker ezrin is elevated at the back of the cell (*129, 130*). These observations suggest that membrane-cortex attachment could play a role in membrane deformations. Meanwhile, we quantitatively determined the critical conditions for this onset event and found a linear dependence between these thresholds and linker density. The second prediction of our model is that the glycocalyx alone can drive the production of extracellular spherical microvesicles under the condition that a threshold grafting density is reached. This prediction is supported by experimental evidence showing that the removal of the glycocalyx leads to a significant reduction in the release of vesicles (*53*). This might also partially have implications for understanding microvesicle generation in many cancer-cell types. For instance, tumor cells with high mucin expression have a propensity to generate a high number of microvesicles (*131*). Moreover, the predicted dependence of critical grafting density on various system parameters for the successful generation of extracellular spherical microvesicles may provide guidance on how to adjust control parameters to fine-tune membrane geometries in experiments. In the end, our model also predicts that a heterogeneous collection of EV population can be produced through the regulation of glycocalyx properties. This prediction is supported by the experimental observations made by Shurer et al. (*53*), and the underlying mechanisms may help explain why EVs are heterogeneous structures spanning a wide range of sizes. We expect that the findings in this work can serve as a better understanding of the role of membrane-cortex attachment and individual glycocalyx properties in regulating membrane shapes and provide deeper insights into how extracellular vesiculation occurs under various biophysical conditions.

To investigate the effects of membrane-cortex adhesion and the glycocalyx on EV formation and to incorporate these factors into our theoretical model, we made several assumptions and simplifications in the present work. First, we approximated the shape of the membrane as a family of spherical caps, which is a reasonable simplification since Shurer et al. (*53*) reported that bulky brush-like glycocalyx polymers are sufficient to induce spherical-shaped membranes (referred to as blebs). Second, the model does not include the electrostatic effects because we treated the glycocalyx as an uncharged polymer brush network. Third, we neglected the spatial heterogeneity of the distribution of the glycocalyx polymers on the membrane. Fourth, beyond the local breakage of bonds between the cortex and the membrane, diffusion and binding and unbinding of the protein linker may also affect glycocalyx-mediated membrane architectures within the cell (*57, 64*). Here, we neglected the diffusion and unbinding kinetics of the protein linker in the cortex. In addition, the energy associated with membrane-cortex adhesion was calculated using the spring constant of the linker and its displacement. Future modeling and experimental studies could incorporate the diffusion of glycocalyx polymers, changes in the bulk physical properties of the glycocalyx, charged polymers, as well as the effects of linker binding and unbinding in the actin cortex to better capture the biological complexity of cell membranes.

Combining polymer physics-based theory and Helfrich membrane theory, we proposed a theoretical model to investigate the mechanisms by which glycocalyx polymers and membrane-cortex adhesion influence membrane shape remodeling. Specifically, a minimal model allows us to quantify the roles of glycocalyx, membrane-cortex adhesion, grafting area, line tension, and spontaneous curvature in the formation of EVs. The physics behind the transition between different deformation states comes from the competition among the energy associated with the glycocalyx, the elastic energy (consisting of bending energy and tension energy), and membrane-cortex adhesion energy. The energy associated with the glycocalyx drive membrane deformation at the cost of the bending energy due to local membrane curvature formation, of stretching energy due to lateral membrane tension, and of adhesion energy due to the attachment between the membrane and the cortex. Line tension and spontaneous curvature are also driving factors assisting membrane deformation.

In conclusion, based on the developed theoretical framework, we study the EV formation regulated by glycocalyx and membrane-cortex adhesion and find that the presence of glycocalyx is able to initiate membrane deformation and regulate EV size distribution. Such membrane shape remodeling behaviors induced by glycocalyx and membrane-cortex adhesion provide a deeper understanding of EV secretion. EV formation is tunable by glycocalyx, cortex-membrane adhesion, grafting area, line tension, and spontaneous curvature, which can be characterized by a phase diagram in the two-parameter space among these factors. We identify that the degree of membrane deformation can be increased upon regulating many parameters, such as increasing glycocalyx grafting density and length, reducing cortex-membrane adhesion strength by tuning the linker density and stiffness, and increasing line tension and spontaneous curvature.

## Acknowledgments

This work was supported by NIH R01GM132106 and Office of Naval Research N00014-20-1-2469 to P.R. We would like to thank Dr. Emmet Francis for his thoughtful feedback.

## Supporting Information Text (SI)

### 1 Derivation of the energy contribution associated with the glycocalyx

Here we derive Eq. (20) of the main text. Notably, the crowding of large glycosylated proteins appears to regulate the shape of the underlying bilayer plasma membrane (*53*). According to polymer physics, two regimes can be defined depending on the grafting density of glycocalyx polymers on the cell membrane, which are the mushroom-like regime and the brush-like regime. Shurer et al. (*53*) have reported that the mucins are in the coiled mushroom state in the case of low-density glycocalyx polymers. We focus on densely grafted regions of the membrane and therefore model the glycocalyx in the brush regime.

**Figure S1.**
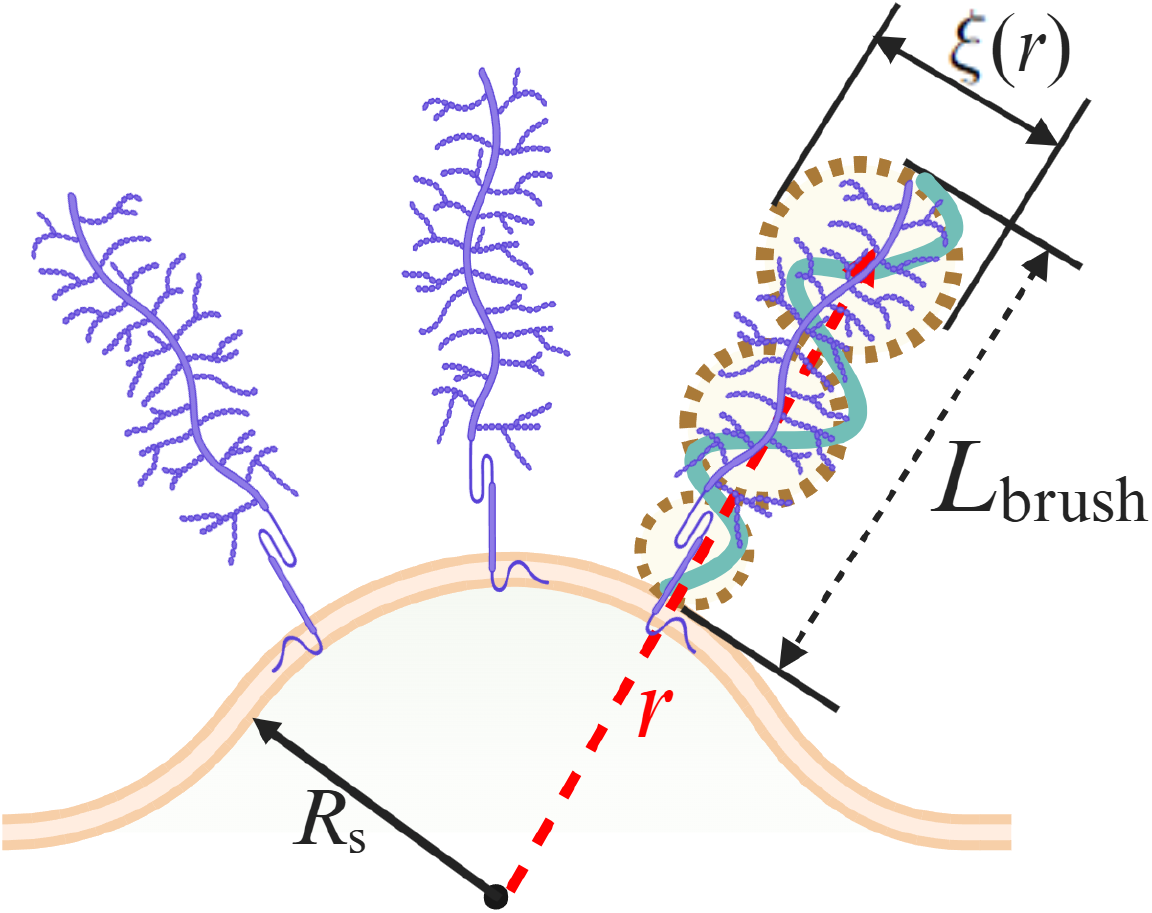
Schematic of a layer of brush-like polymers attached on a spherical cap membrane surface with radius *R*_s_. At a given position *r*, the coarse grained blob (dashed black circle) size in a polymer brush is *ξ*(*r*), the thickness of the brush is *L*_brush_.

Here, we consider a layer of glycocalyx grafted on a membrane of radius *R*_s_ in a brush-like structure with thickness *L*_brush_, as shown in Fig. S1. From the viewpoint of coarse graining, the polymer brush is envisioned as an array of blobs. The size of each blob, *ξ*, at a given position *r* equals the square root of the local area per chain *s*(*r*), where *r* = *x* + *R*_s_ is the radial distance which is defined from the center of the spherical surface, and in which *x* is the distance from the membrane surface. Thus, the blob size 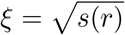 grows as a function of *r*, and the grafting density of glycocalyx polymer brush on the membrane surface can be obtained as *ρ* = 1*/ξ*^2^. Assuming that the layer of glycopolymers are extended non-uniformly but equally in the height, *L*_brush_, then the area per chain at distance *x* from the membrane surface is given by (*106*)

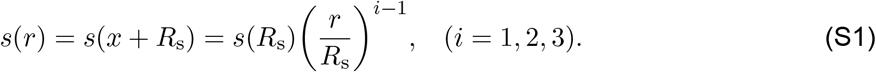

The index *i* = 1, 2, 3 indicates planar, cylindrical, and spherical shaped membranes, respectively. When the membrane is bent, the changes of polymer configuration give rise to the local extension of the polymer chain. Here the local chain extension at a height *r* is characterized by d*r/*d*n*, where the variable *n* denotes the current monomer. This local extension is related to local density profile of monomers *c*_p_(*r*) as (*106*)

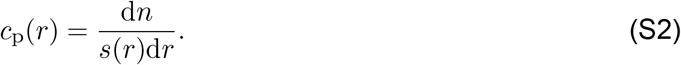

Then, the thickness of the brush, *L*_brush_, is found from the conservation condition (the constraint of conservation of the total number of monomers *N*) (*106*),

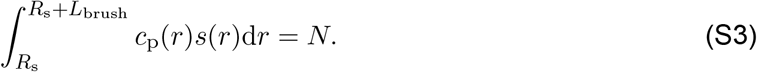

As a result, in the brush regime, since electrostatic effects are excluded, the energy contribution originating from glycocalyx polymers includes two terms (*53, 106*): the elastic energy of the polymer chain (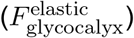) and the free energy caused by the excluded volume interactions of polymer monomers (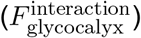). Based on the hypotheses in the main text, according to Ref. (*106*), the elastic energy per chain in the brush can be represented as

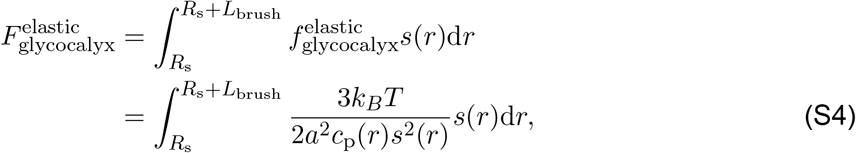

where *a* is the monomer length and 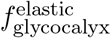 is the elastic energy density of the polymer chain. Within the mean-field approximation, the energy density of the excluded volume interactions (van der Waals interactions) between monomers can be modeled in terms of the virial expansion

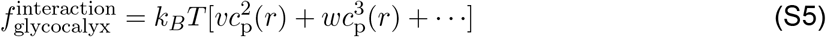

where *v* and *w* are the second and third virial coefficient, respectively. As a result, the excluded volume interactions between monomers per chain in the brush is given by

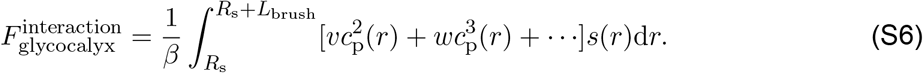

Moving forward, the cubic and higher terms are neglected. Therefore, the sum of Eq. (S4) and Eq. (S6) yields the energy contribution of glycocalyx polymers

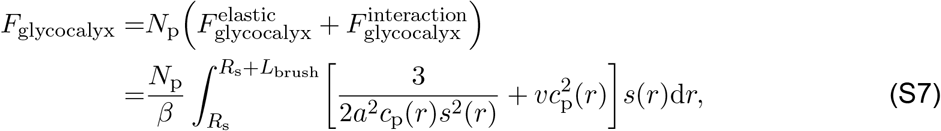

where *N*_p_ = *ρA*_0_ is the number of polymer chain grafted on the membrane. Here *A*_0_ stands for the membrane patch area covered by glycocalyx polymers and *ρ* is the density.

In order to calculate the free energy, we need to further determine the local concentration of monomers *c*_p_(*r*) and the brush thickness *L*_brush_. On a planar membrane surface, the planar brush area per chain, *s*(*r*), is constant, i.e., *s*(*r*) = *s*. Minimizing the free energy of a planar brush *F*_glycocalyx_ with respect to *c*_p_ by using 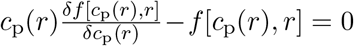 leads to the equilibrium polymer concentration 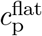 (see Ref. (*106*) for detailed steps)

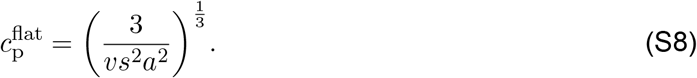

Here, 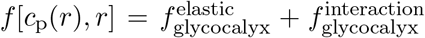. Using the conservation condition defined by Eq. (S3) yields the brush thickness 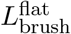 see Ref. (*106*) for detailed steps)

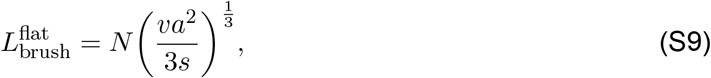

where we can see that the thickness of polymer brush is proportional to the total number of effective monomers *N*. Hereafter, in our model, we use the number of effective monomers, *N*, to capture the length of the polymer. Note that the relationship for 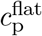 and 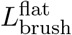 are special cases of electrically neutral brushes as described in (*106*). Substituting Eq. (S8) and Eq. (S9) into Eq. (S7) yields the energy contribution of glycocalyx polymers on a planar membrane surface

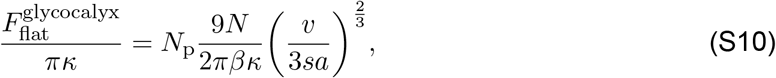

which corresponds to the case *η* = 0 in Eq. (20) in the main text. Thus, even for a flat membrane, the energy contribution by the glycocalyx is directly proportional to the extent of grafting *N*_p_ and the length of the polymer brush *N*.

On a spherical membrane surface, the corresponding monomer density profile *c*_p_(*r*) and brush thickness *L*_brush_ are, respectively, expressed as (see Ref. (*106*) for a detailed calculation)

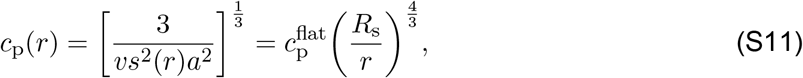

and

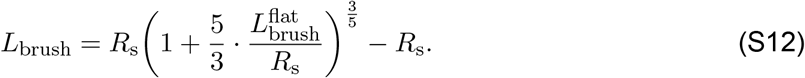

Similarly, substituting Eq. (S11) and Eq. (S12) into Eq. (S7) leads to the energy contribution of glycocalyx polymers on a spherical membrane surface as

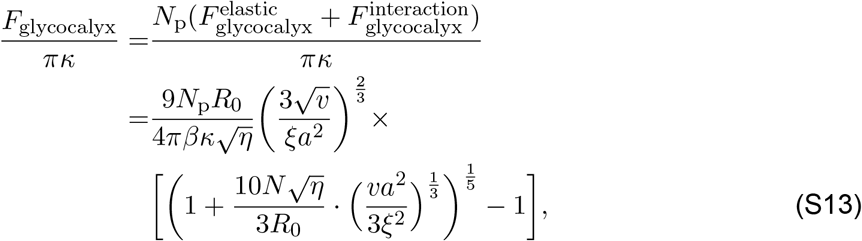

where we use 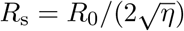.

### 2 Derivation of the energy associated with membrane-cortex adhesion

Here we derive Eq. (22) of the main text. To estimate the energy associated with membrane-cortex adhesion, we assume that the membrane-cortex adhesion energy is quadratic with respect to the membrane-cortex linker deformation and is thus given by

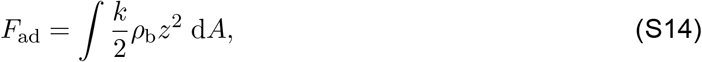

where *k* is the elastic stiffness (spring constant) of each membrane-cortex linker, *ρ*_b_ is the local density of bound spring-like linkers, and *z* is the stretching length of linkers. For simplicity, we further assume that (i) the density of bound spring-like linkers equals the density of available linkers *ρ*_0_, i.e., *ρ*_b_ = *ρ*_0_; (ii) the adhesion bond breaks as the membrane-cortex adhesion bond reaches its maximum length *l*_c_.

When the membrane deformation is small, the maximum height of the deformed membrane is smaller than the maximum extension length of linkers (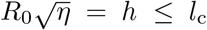 which corresponds to *η ≤*(*l*_c_*/R*_0_)^2^, see Fig. 2(b) in the main text), the membrane-cortex adhesion energy can be calculated as

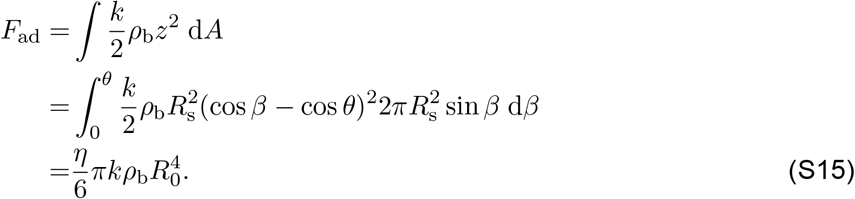

where cos *θ* = 1 *−* 2*η* with *θ* being the opening angle of the spherical cap, and *β* is the integral variable.

If the membrane is deformed into a spherical cap with its maximum height larger than the maximum extension length of the linkers, that is, *h > l*_c_ (see Fig. 2(c) in the main text). Then, it means that part of the linkers are broken (see the regime above the dashed line in Fig. 2(c) in the main text) and some of the linkers are still attached to the membrane but with some degree of extension (see the regime below the dashed line in Fig. 2(c) in the main text) for the deformed membrane. Thereby, we can treat the membrane as two parts, one with the bond linkers are broken, and the other one with the linkers binding on it. As a result, in the case of *h > l*_c_, the membrane-cortex adhesion energy consists of two parts, the first one is the part with linker broken, and the second one is the detached part with the linkers still attached but extended.

When the membrane is detached from the cortex, the total number of broken adhesion bonds can be approximated by *N*_b_ = *ρ*_b_*A*_d_(*η*), where *A*_d_(*η*) is the area of the detached membrane, which represents the area where the adhesion between the membrane and the cortex has been disrupted, as shown in Fig. 2(c) in the main text. Specifically, if we assume the detachment between the membrane and the cortex occurs once their separation reaches *l*_c_, then *A*_d_(*η*) can be shown to take the form

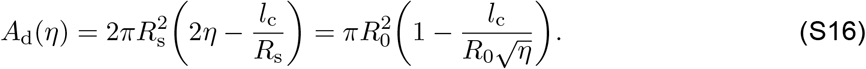

Therefore, the energy contribution from the first part can be obtained as

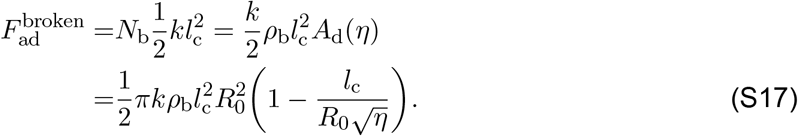

The energy contribution from the second part can be obtained as

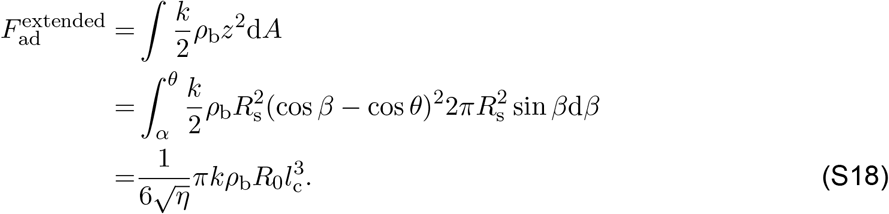

Consequently, the membrane-cortex adhesion energy can be expressed as

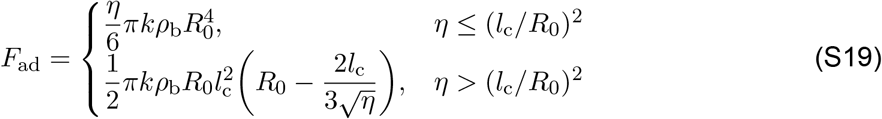

which coincides with Eq. (22) of the main text.

### 3 The effects of spontaneous curvature and membrane tension on system stability

To investigate the effects of spontaneous curvature and membrane tension on system stability, the dispersion relations for different values of spontaneous curvature and membrane tension are plotted, as shown in Fig. S2. Figure S2(a) shows that the instability appears when the spontaneous curvature reaches 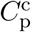, and the eigenvalue becomes unstable in a range of wavevectors *q*_*−*_ *< q < q*_+_ when *C*_p_ is smaller than 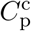. Moreover, *q*_*−*_ can vanish if *C*_p_ is small enough (pink curve). In Fig. S2(b), we plot the dependence of the unstable wavenumbers on the spontaneous curvature, where we also observe two types of instabilities similar to those of Fig. 3(d). In Fig. S2(c), we plot the dispersion relations for a set of fixed membrane tensions, where the curves show a trend similar to that in Fig. 3(a). In the end, the stability phase diagram of the *q− σ* plane demonstrates a cT-type instability (Fig. S2(d)).

**Figure S2.**
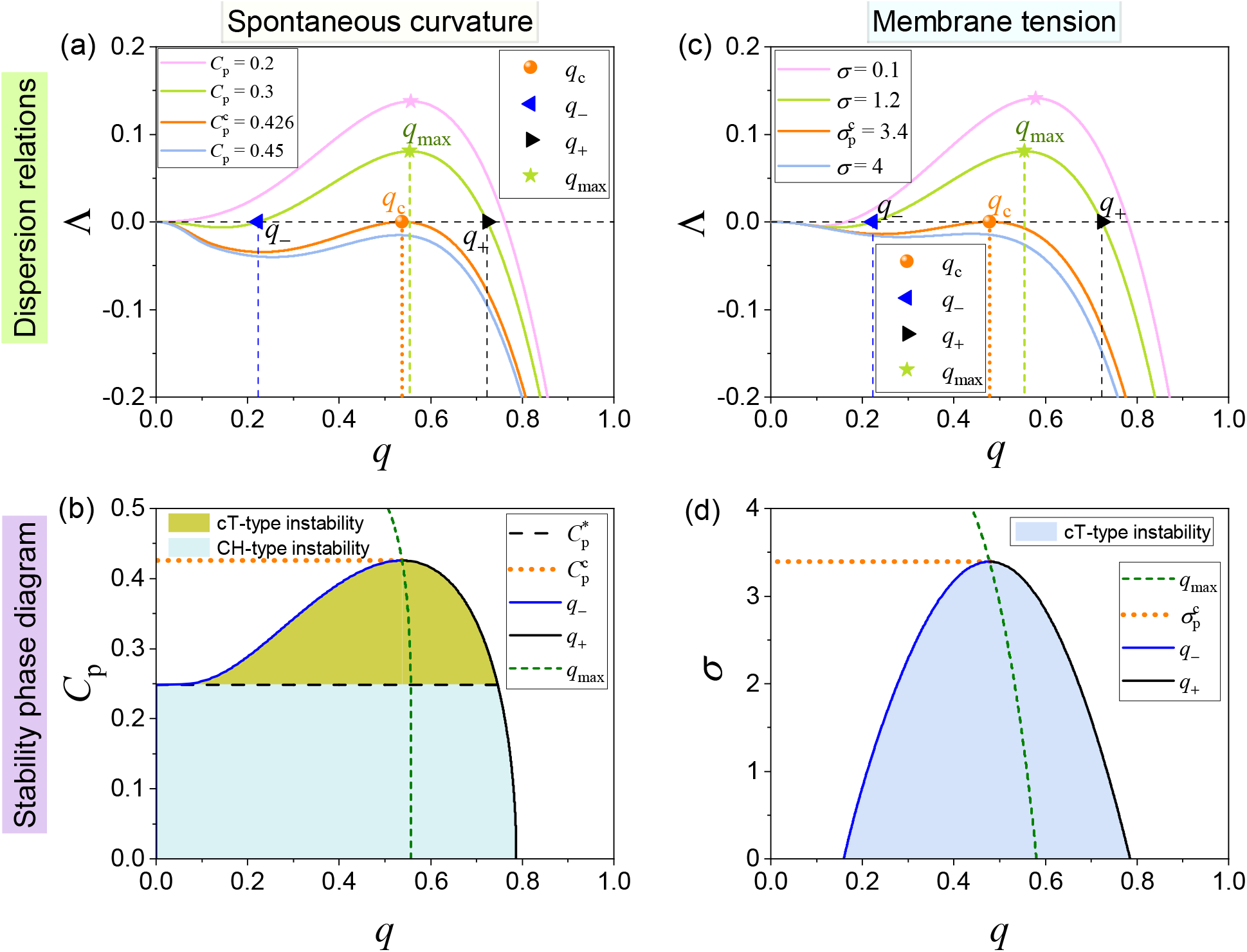
Linear stability analysis of the composite system. Plot of the dispersion relation Λ(*q*) for different values of (a) spontaneous curvature and (c) membrane tension. The dependence of the unstable wavenumbers on the (b) spontaneous curvature and (d) membrane tension.

### 4 Effects of spontaneous curvature and line tension on EV formation

To intuitively observe the influence of spontaneous curvature and line tension on EV formation, a phase diagram on the (*λ − c*_0_) plane is constructed, as shown in Fig. S3. For zero line tension and spontaneous curvature, the cell membrane deformation is in a shallow spherical cap state. By increasing line tension and spontaneous curvature to certain values, the transition from an inflated state to a full EV state can be observed, indicating a first-order transition. This suggests that line tension *λ* and spontaneous curvature *c*_0_ play a positive role in the formation of EVs.

**Figure S3.**
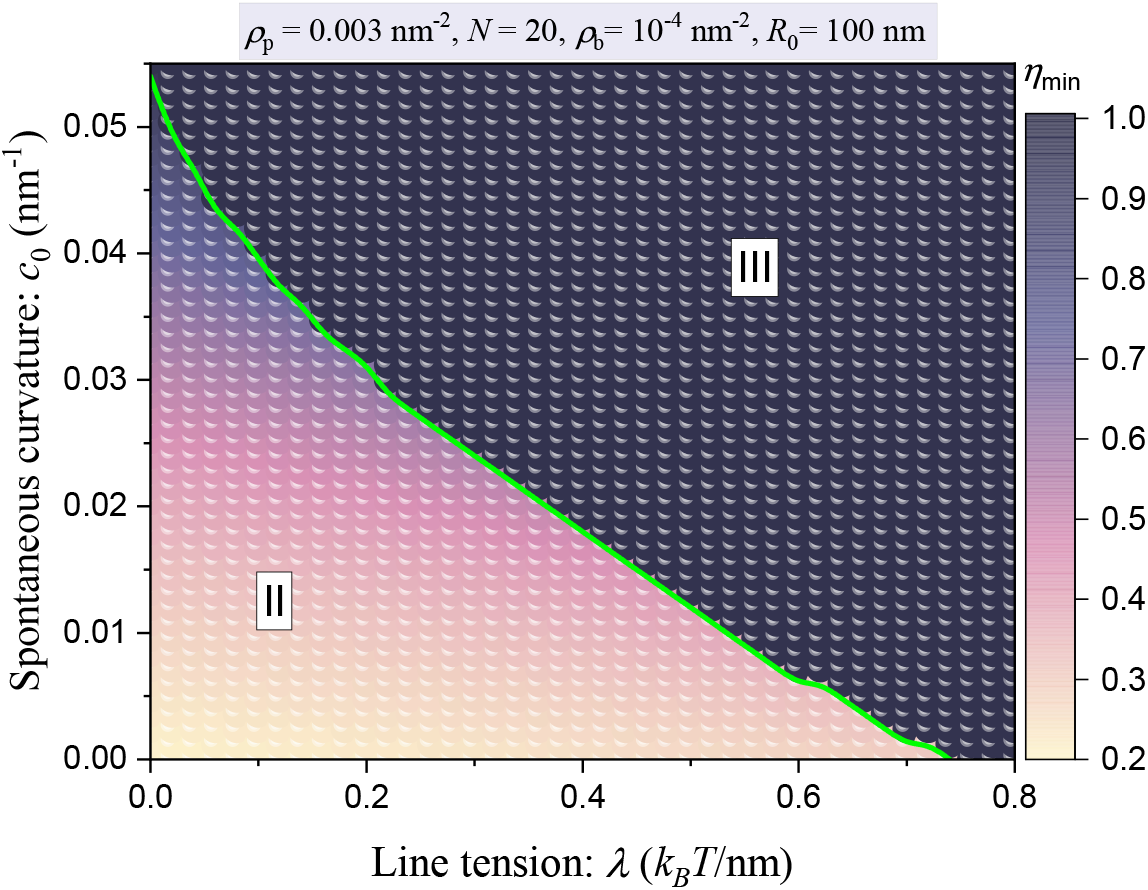
Contour plot of optimal shape parameter *η*_min_ as a function of line tension *λ* and spontaneous curvature *c*_0_, where the color bar represents the magnitude of the optimal shape parameter.

### 5 Phase diagrams on the (*R*_0_ *− ρ*_b_) plane

To gain more insight into the effects of glycocalyx properties on EV formation, two phase diagrams are constructed on the *R*_0_ *− ρ*_b_ plane for different values of glycocalyx grafting density and glycopolymer length, as shown in Fig. S4. When comparing Fig. S4(a) with Fig. 6(e), we observe that an elevate of glycocalyx grafting density leads to a narrow regime I and shifts the boundary curve to the right (see the cyan curve). Furthermore, the comparison of regime I between Fig. S4(b) and Fig. 6(e) illustrates that the flat membrane regime also becomes narrower, and the boundary curve (magenta curve) that separates regime I from regime II shifts to the right in the case of longer glycopolymer length. These indicate that a higher threshold of the linker density and glycocalyx grafting area is required to prohibit the formation of EVs in the presence of a dense and long glycocalyx cell membrane. This also further validates the prediction that the glycocalyx promotes the formation of EVs.

**Figure S4.**
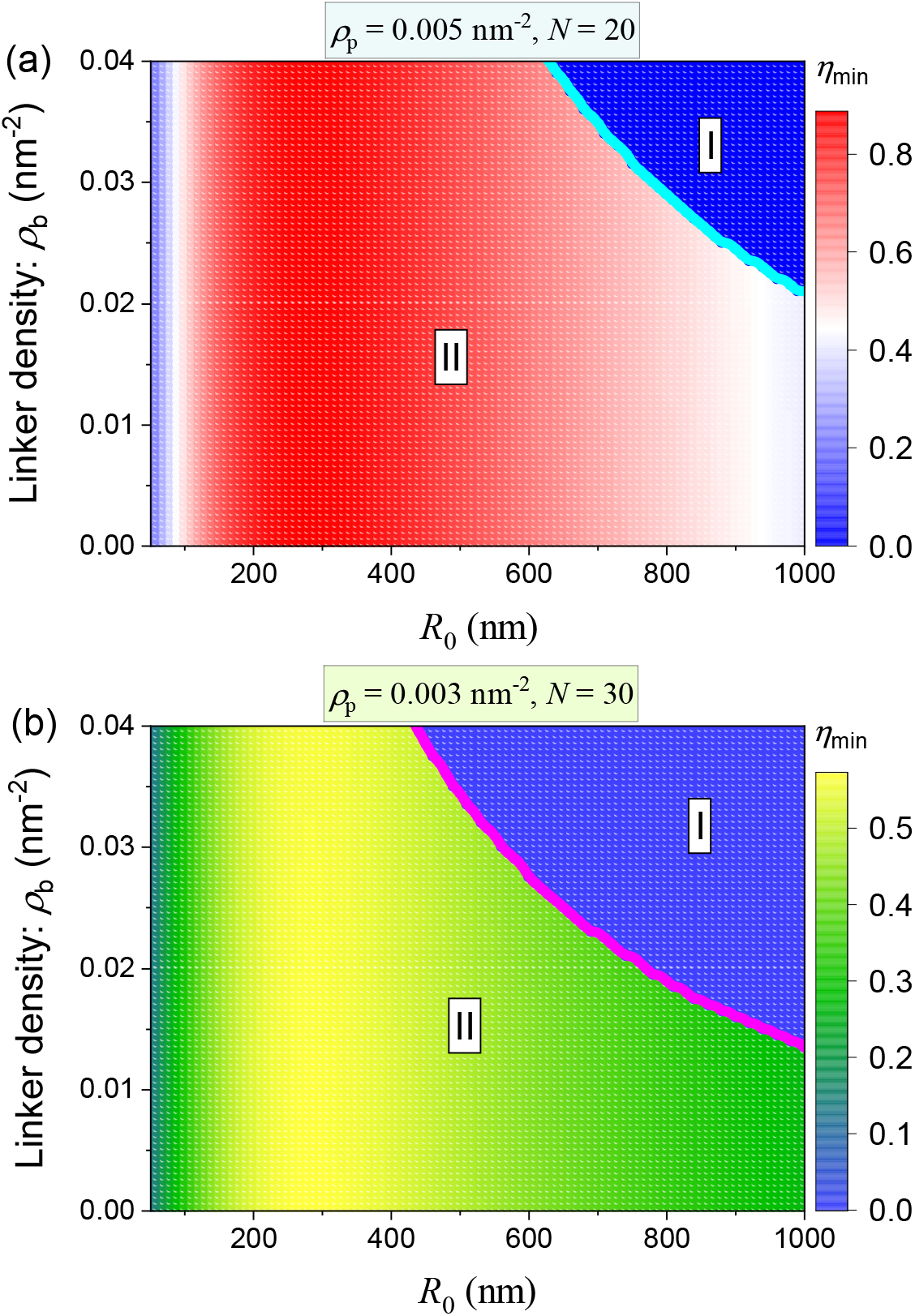
Contour plot of optimal shape parameter *η*_min_ as a function of patch radius *R*_0_ and linker density *ρ*_b_ for the grafting density and length of the glycocalyx are fixed as (a) *ρ*_p_ = 0.005 nm^*−*2^ and *N* = 20, and (b) *ρ*_p_ = 0.003 nm^*−*2^ and *N* = 30, where the color bar represents the magnitude of the optimal shape parameter.

## References

1. D. K. Jeppesen, Q. Zhang, J. L. Franklin, R. J. Coffey, Trends Cell Biol. 33, 667–681, ISSN: 0962-8924, DOI 10.1016/j.tcb.2023.01.002 (2023).

2. D. K. Jeppesen et al., Cell 177, 428–445.e18, ISSN: 0092-8674, DOI 10.1016/j.cell.2019.02.029 (2019).

3. M. A. Kumar et al., Sig. Transduct. Target. Ther. 9, 27, ISSN: 2095-9907, DOI 10.1038/s41392-024-01735-1 (Feb. 2024).

4. J. A. Welsh et al., J. Extracell. Vesicles 13, e12404, DOI 10.1002/jev2.12404 (2024).

5. M. Sheta, E. A. Taha, Y. Lu, T. Eguchi, Biol. 12, 110, ISSN: 2079-7737, DOI 10.3390/biology12010110 (2023).

6. T. Ahmadzada et al., Crit. Rev. Oncol. Hematol. 150, 102949, ISSN: 1040-8428, DOI 10.1016/j.critrevonc.2020.102949 (2020).

7. A. Becker et al., Cancer Cell 30, 836–848, ISSN: 1535-6108, DOI 10.1016/j.ccell.2016.10.009 (2016).

8. L. A. Hargett, N. N. Bauer, Pulm. Circ. 3, 329–340, DOI 10.4103/2045-8932.114760 (2013).

9. K. Segawa, J. Suzuki, S. Nagata, Proc. Natl. Acad. Sci. U. S. A. 108, 19246–19251, DOI 10.1073/pnas.1114799108 (2011).

10. J. D. Arroyo et al., Proc. Natl. Acad. Sci. U. S. A. 108, 5003–5008, DOI 10.1073/pnas.1019055108 (2011).

11. K. C. Vickers, B. T. Palmisano, B. M. Shoucri, R. D. Shamburek, A. T. Remaley, Nat. Cell Biol. 13, 423–U182, ISSN: 1465-7392, DOI 10.1038/ncb2210 (Apr. 2011).

12. H. Zhao et al., Biochim. Biophys. Acta, Rev. Cancer 1869, 64–77, ISSN: 0304-419X, DOI 10.1016/j.bbcan.2017.11.005 (2018).

13. D. E. Reynolds et al., J. Ex. Bio. 2, e89, DOI 10.1002/jex2.89(2023).

14. S. Muraoka et al., Alzheimer’s Dement. 16, 896–907, DOI 10.1002/alz.12089 (2020).

15. C. Jiang et al., Mov. Disord. 36, 2663–2669, DOI 10.1002/mds.28591 (2021).

16. M.O. Gonçalves et al., in Extracellular Vesicles from Basic Research to Clinical Applications, ed. by A. C. Torrecilhas (Academic Press, 2024), vol. 94, pp. 1–31, DOI 10.1016/bs.ctm.2024.06.008.

17. K. Abhange et al., Bioact. Mater. 6, 3705–3743, ISSN: 2452-199X, DOI 10.1016/j.bioactmat.2021.03.015 (2021).

18. K. D. Brett et al., Sci. Rep. 9, 13320, ISSN: 2045-2322, DOI 10.1038/s41598-019-49924-1 (Sept. 2019).

19. J. H. W. Distler et al., Arthritis & Rheumatism 52, 3337–3348, DOI 10.1002/art.21350 (2005).

20. D. Ruhela et al., Brain Behav. Immun. 87, 465–472, ISSN: 0889-1591, DOI 10.1016/j.bbi.2020.01.017 (2020).

21. D. Ruhela, V. M. Bhopale, S. Kalakonda, S. R. Thom, Brain Behav. Immun. Health 18, 100398, ISSN: 2666-3546, DOI 10.1016/j.bbih.2021.100398 (2021).

22. C. Ciardiello, R. Migliorino, A. Leone, A. Budillon, Cytokine Growth Factor Rev. 51, 69–74, ISSN: 1359-6101, DOI 10.1016/j.cytogfr.2019.12.007 (2020).

23. Z. Varga et al., Colloids Surf. B Biointerfaces 192, 111053, ISSN: 0927-7765, DOI 10.1016/j.colsurfb.2020.111053 (2020).

24. M. F. Baietti et al., Nat. Cell Biol. 14, 677–685, ISSN: 1465-7392, DOI 10.1038/ncb2502 (July 2012).

25. M. Ostrowski et al., Nat. Cell Biol. 12, 19–U61, ISSN: 1465-7392, DOI 10.1038/ncb2000 (Jan. 2010).

26. F. K. Fordjour, C. Guo, Y. Ai, G. G. Daaboul, S. J. Gould, J. Biol. Chem. 298, 102394, ISSN: 0021-9258, DOI 10.1016/j.jbc.2022.102394 (2022).

27. M. Mathieu et al., Nat. Commun. 12, 4389, ISSN: 2041-1723, DOI 10.1038/s41467-021-24384-2 (July 2021).

28. T. Matsui, F. Osaki, S. Hiragi, Y. Sakamaki, M. Fukuda, EMBO Rep. 22, e51475, DOI 10.15252/embr.202051475 (2021).

29. J. Kowal et al., Proc. Natl. Acad. Sci. U. S. A. 113, E968–E977, DOI 10.1073/pnas.1521230113 (2016).

30. M. Dieudé et al., Sci. Transl. Med. 7, 318ra200–318ra200, DOI 10.1126/scitranslmed.aac9816 (2015).

31. M. Hristov, W. Erl, S. Linder, P. C. Weber, Blood 104, 2761–2766, ISSN: 0006-4971, DOI 10.1182/blood-2003-10-3614 (Nov. 2004).

32. L. Ma et al., Cell Res. 25, 24–38, ISSN: 1001-0602, DOI 10.1038/cr.2014.135 (Jan. 2015).

33. X. Zhao et al., Cell Discov. 5, 27, ISSN: 2056-5968, DOI 10.1038/s41421-019-0093-y (May 2019).

34. Y. Huang et al., Nat. Cell Biol. 21, 991–1002, ISSN: 1465-7392, DOI 10.1038/s41556-019-0367-5 (Aug. 2019).

35. V. R. Minciacchi et al., Cancer Res. 77, 2306–2317, ISSN: 0008-5472, DOI 10.1158/0008-5472.CAN-16-2942 (Apr. 2017).

36. D. Di Vizio et al., Am. J. Pathol. 181, 1573–1584, ISSN: 0002-9440, DOI 10.1016/j.ajpath.2012.07.030 (2012).

37. A. Callan-Jones, P. Bassereau, Curr. Opin. Solid State Mater. Sci. 17, 143–150, DOI 10.1016/j.cossms.2013.08.004 (2013).

38. M. M. Kozlov, J. W. Taraska, Nat. Rev. Mol. Cell Biol. 24, 63–78, DOI 10.1038/s41580-022-00511-9 (2023).

39. J. Derganc, A. Čopič, Biochim. Biophys. Acta 1858, 1152–1159, DOI 10.1016/j.bbamem.2016.03.009 (2016).

40. J. C. Stachowiak et al., Nat. Cell Biol. 14, 944+, DOI 10.1038/ncb2561 (2012).

41. D. J. Busch et al., Nat. Commun. 6, 7875, DOI 10.1038/ncomms8875 (2015).

42. S. Liese, A. Carlson, Biophys. J. 120, 2482–2489, DOI 10.1016/j.bpj.2021.04.029 (2021).

43. V. T. Ruhoff, G. Moreno-Pescador, W. Pezeshkian, P. M. Bendix, Biochem. Soc. Trans. 50, 1257–1267, DOI 10.1042/BST20210883 (2022).

44. H. Alimohamadi, P. Rangamani, Biomolecules 8, 120, DOI 10.3390/biom8040120 (2018).

45. N. Walani, J. Torres, A. Agrawal, Proc. Natl. Acad. Sci. U. S. A. 112, E1423–E1432, DOI 10.1073/pnas.1418491112 (2015).

46. J. E. Hassinger, G. Oster, D. G. Drubin, P. Rangamani, Proc. Natl. Acad. Sci. U. S. A. 114, E1118–E1127, DOI 10.1073/pnas.1617705114 (2017).

47. R. Ma, J. Berro, Biophys. J. 120, 1625–1640, DOI 10.1016/j.bpj.2021.02.033 (2021).

48. K. Xiao, C.-X. Wu, R. Ma, Phys. Rev. Res. 5, 023176, DOI 10.1103/PhysRevResearch.5.023176 (2 June 2023).

49. B. Alberts et al., presented at the (Garland Science, New York, 2015).

50. J. C.-H. Kuo, J. G. Gandhi, R. N. Zia, M. J. Paszek, Nat. Phys. 14, 658–669, DOI 10.1038/s41567-018-0186-9 (2018).

51. S. Weinbaum, J. M. Tarbell, E. R. Damiano, Annu. Rev. Biomed. Eng. 9, 121–167, DOI 10.1146/annurev.bioeng.9.060906.151959 (2007).

52. L. Möckl, Front. Cell Dev. Biol. 8, 253, DOI 10.3389/fcell.2020.00253 (2020).

53. C. R. Shurer et al., Cell 177, 1757–1770.e21, DOI 10.1016/j.cell.2019.04.017 (2019).

54. J. C.-H. Kuo, M. J. Paszek, Annu. Rev. Cell Dev. Biol. 37, 257–283, DOI 10.1146/annurev-cellbio-120219-054401 (2021).

55. S. Gollapudi et al., Proc. Natl. Acad. Sci. U. S. A. 120, e2215815120, DOI 10.1073/pnas.2215815120 (2023).

56. R. Alert, J. Casademunt, Phys. Rev. Lett. 116, 068101, DOI 10.1103/PhysRevLett.116.068101 (6 Feb. 2016).

57. R. Alert, J. Casademunt, J. Brugués, P. Sens, Biophys. J. 108, 1878–1886, DOI 10.1016/j.bpj.2015.02.027 (2015).

58. F. Y. Lim, K.-H. Chiam, L. Mahadevan, Europhys. Lett. 100, 28004, DOI 10.1209/0295-5075/100/28004 (2012).

59. W. Strychalski, R. D. Guy, Math. Med. Biol. 30, 115–130, DOI 10.1093/imammb/dqr030 (2012).

60. W. Strychalski, R. D. Guy, Biophys. J. 110, 1168–1179, DOI 10.1016/j.bpj.2016.01.012 (2016).

61. E. K. Paluch, E. Raz, Curr. Opin. Cell Biol. 25, 582–590, DOI 10.1016/j.ceb.2013.05.005 (2013).

62. G. Charras, E. Paluch, Nat. Rev. Mol. Cell Biol. 9, 730–736, DOI 10.1038/nrm2453 (2008).

63. R. Merkel et al., Biophys. J. 79, 707–719, DOI 10.1016/S0006-3495(00)76329-6 (2000).

64. E. S. Welf et al., Dev. Cell 55, 723–736.e8, DOI 10.1016/j.devcel.2020.11.024 (2020).

65. J. Dai, M. P. Sheetz, Biophys. J. 77, 3363–3370, DOI 10.1016/S0006-3495(99)77168-7 (1999).

66. A. Diz-Muñoz et al., PLoS Biol. 8, 1–12, DOI 10.1371/journal.pbio.1000544 (2010).

67. A. Paraschiv et al., Biophys. J. 120, 598–606, DOI 10.1016/j.bpj.2020.12.028 (2021).

68. M. Sheetz, Nat. Rev. Mol. Cell Biol. 2, 392–396, DOI 10.1038/35073095 (2001).

69. F.-C. Tsai, G. Guérin, J. Pernier, P. Bassereau, Eur. J. Cell Biol. 103, 151402, ISSN: 0171-9335, DOI 10.1016/j.ejcb.2024.151402 (2024).

70. M. Tsujioka et al., Proc. Natl. Acad. Sci. U. S. A. 109, 12992–12997, DOI 10.1073/pnas.1208296109 (2012).

71. S. Tsukita, S. Yonemura, J. Biol. Chem. 274, 34507–34510, DOI 10.1074/jbc.274.49.34507 (1999).

72. R. G. Fehon, A. I. McClatchey, A. Bretscher, Nat. Rev. Mol. Cell Biol. 11, 276–287, DOI 10.1038/nrm2866 (2010).

73. E. Korkmazhan, A. R. Dunn, Sci. Adv. 8, eabo2779, DOI 10.1126/sciadv.abo2779 (2022).

74. K. Xiao, S. Park, J. C. Stachowiak, P. Rangamani, Proc. Natl. Acad. Sci. U. S. A. 122, e2418357122, DOI 10.1073/pnas.2418357122 (2025).

75. K. Xiao, P. Rangamani, Biophys. J. 124, 1631–1642, DOI 10.1016/j.bpj.2025.04.006 (2025).

76. C. L. Hattrup, S. J. Gendler, Annu. Rev. Physiol. 70, 431–457, DOI 10.1146/annurev.physiol.70.113006.100659 (2008).

77. Y. Jung et al., Proc. Natl. Acad. Sci. U. S. A. 113, E5916–E5924, DOI 10.1073/pnas.1605399113 (2016).

78. G. Kesavan et al., Cell 139, 791–801, DOI 10.1016/j.cell.2009.08.049 (2009).

79. M. Kesimer et al., Mucosal Immunol. 6, 379–392, DOI 10.1038/mi.2012.81 (2013).

80. S. Makabe, T. Naguro, T. Stallone, Microsc. Res. Tech. 69, 436–449, DOI 10.1002/jemt.20303 (2006).

81. S. P. Evanko, M. I. Tammi, R. H. Tammi, T. N. Wight, Adv. Drug Delivery Rev. 59, 1351– 1365, DOI 10.1016/j.addr.2007.08.008 (2007).

82. B. Button et al., Science 337, 937–941, DOI 10.1126/science.1223012 (2012).

83. J. Richard Bennett et al., J. Histochem. Cytochem. 49, 67–77, DOI 10.1177/002215540104900107 (2001).

84. V. Koistinen et al., Exp. Cell Res. 337, 179–191, DOI 10.1016/j.yexcr.2015.06.016 (2015).

85. C. Hiergeist, R. Lipowsky, J. Phys. II France 6, 1465–1481, DOI 10.1051/jp2:1996142 (1996).

86. R. Lipowsky, Europhys. Lett. 30, 197, DOI 10.1209/0295-5075/30/4/002 (1995).

87. M. Breidenich, R. R. Netz, R. Lipowsky, Europhys. Lett. 49, 431, DOI 10.1209/epl/i2000-00167-2 (2000).

88. T. Bickel, C. Jeppesen, C. Marques, Eur. Phys. J. E 4, 33–43, DOI 10.1007/s101890170140 (2001).

89. Y. W. Kim, W. Sung, Phys. Rev. E 63, 041910, DOI 10.1103/PhysRevE.63.041910 (2001).

90. F. Campelo, A. Hernández-Machado, Phys. Rev. Lett. 99, 088101, DOI 10.1103/PhysRevLett.99.088101 (2007).

91. F. Campelo, A. Hernández–Machado, Phys. Rev. Lett. 100, 158103, DOI 10.1103/PhysRevLett.100.158103 (2008).

92. M. Werner, J. U. Sommer, Eur. Phys. J. E 31, 383–392, DOI 10.1140/epje/i2010-10576-4 (2010).

93. S. Kutti Kandy, R. Radhakrishnan, Biophys. J. 121, 3674–3683, DOI 10.1016/j.bpj.2022.05.031 (2022).

94. G. Charras, J. Microsc. 231, 466–478, DOI 10.1111/j.1365-2818.2008.02059.x (2008).

95. J.-Y. Tinevez et al., Proc. Natl. Acad. Sci. U. S. A. 106, 18581–18586, DOI 10.1073/pnas.0903353106 (2009).

96. A. Mahapatra, S. A. Malingen, P. Rangamani, biorxiv, DOI 10.1101/2024.02.07.579325 (2024).

97. P. Canham, Journal of Theoretical Biology 26, 61–81, ISSN: 0022-5193, DOI 10.1016/S0022-5193(70)80032-7 (1970).

98. W. Helfrich, Z. Naturforsch. C 28, 693–703, DOI 10.1515/znc-1973-11-1209 (1973).

99. P. J. Flory, J. Chem. Phys. 9, 660–660, ISSN: 0021-9606, DOI 10.1063/1.1750971 (Aug. 1941).

100. M. L. Huggins, J. Chem. Phys. 9, 440–440, ISSN: 0021-9606, DOI 10.1063/1.1750930 (May 1941).

101. H. Kramers, Physica 7, 284–304, ISSN: 0031-8914, DOI 10.1016/S0031-8914(40)90098-2 (1940).

102. E. Evans, Annu. Rev. Biophys. 30, 105–128, ISSN: 1936-1238, DOI 10.1146/annurev.biophys.30.1.105 (2001).

103. A. Veksler, N. S. Gov, Biophys. J. 97, 1558–1568, ISSN: 0006-3495, DOI 10.1016/j.bpj.2009.07.008 (2009).

104. R. Shlomovitz, N. S. Gov, Phys. Rev. Lett. 98, 168103, DOI 10.1103/PhysRevLett.98.168103 (16 Apr. 2007).

105. T. Frohoff-Hülsmann, U. Thiele, Phys. Rev. Lett. 131, 107201, DOI 10.1103/PhysRevLett.131.107201 (10 Sept. 2023).

106. E. B. Zhulina, T. M. Birshtein, O. V. Borisov, Eur. Phys. J. E 20, 243–256, DOI 10.1140/epje/i2006-10013-5 (2006).

107. A. Veksler, N. S. Gov, Biophys. J. 93, 3798–3810, ISSN: 0006-3495, DOI 10.1529/biophysj.107.113282 (2007).

108. R. Shlomovitz, N. S. Gov, Phys. Rev. Lett. 98, 168103, DOI 10.1103/PhysRevLett.98.168103 (16 Apr. 2007).

109. A. Winter, Y. Liu, A. Ziepke, G. Dadunashvili, E. Frey, Phys. Rev. E 111, 044405, DOI 10.1103/PhysRevE.111.044405 (4 Apr. 2025).

110. C. Tozzi, N. Walani, M. Arroyo, New J. Phys. 21, 093004, DOI 10.1088/1367-2630/ab3ad6 (Sept. 2019).

111. R. Lipowsky, Biophys. J. 64, 1133–1138, DOI 10.1016/S0006-3495(93)81479-6 (1993).

112. J. Liu, M. Kaksonen, D. G. Drubin, G. Oster, Proc. Natl. Acad. Sci. U. S. A. 103, 10277– 10282, DOI 10.1073/pnas.0601045103 (2006).

113. J.-M. Allain, C. Storm, A. Roux, M. B. Amar, J.-F. Joanny, Phys. Rev. Lett. 93, 158104, DOI 10.1103/PhysRevLett.93.158104 (2004).

114. L. Foret, Eur. Phys. J. E 37, 42, DOI 10.1140/epje/i2014-14042-1 (2014).

115. Z. Shi, T. Baumgart, Nat. Commun. 6, 5974, DOI 10.1038/ncomms6974 (2015).

116. R. Phillips, T. Ursell, P. Wiggins, P. Sens, Nature 459, 379–385, DOI 10.1038/nature08147 (2009).

117. T. Baumgart, S. Hess, W. Webb, Nature 425, 821–824, DOI 10.1038/nature02013 (2003).

118. R. Lipowsky, J. Phys. II France 2, 1825–1840, DOI 10.1051/jp2:1992238 (1992).

119. D. Bracha, E. Karzbrun, G. Shemer, P. A. Pincus, R. H. Bar-Ziv, Proc. Natl. Acad. Sci. U. S. A. 110, 4534–4538, DOI 10.1073/pnas.1220076110 (2013).

120. L. Foret, P. Sens, Proc. Natl. Acad. Sci. U. S. A. 105, 14763–14768, DOI 10.1073/pnas.0801173105 (2008).

121. J. Paturej, S. S. Sheiko, S. Panyukov, M. Rubinstein, Sci. Adv. 2, e1601478, DOI 10.1126/sciadv.1601478 (2016).

122. J. G. Gandhi, D. L. Koch, M. J. Paszek, Biophys. J. 116, 694–708, DOI 10.1016/j.bpj.2018.12.023 (2019).

123. M. C. Cross, P. C. Hohenberg, Rev. Mod. Phys. 65, 851–1112, DOI 10.1103/RevModPhys.65.851 (3 July 1993).

124. M. A. Antonyak et al., Proc. Natl. Acad. Sci. U. S. A. 108, 4852–4857, DOI 10.1073/pnas.1017667108 (2011).

125. T. Wang et al., Proc. Natl. Acad. Sci. U. S. A. 111, E3234–E3242, DOI 10.1073/pnas.1410041111 (2014).

126. M. Goudarzi et al., Dev. Cell 23, 210–218, DOI 10.1016/j.devcel.2012.05.007 (2012).

127. A. Lorentzen, J. Bamber, A. Sadok, I. Elson-Schwab, C. J. Marshall, J. Cell Sci. 124, 1256– 1267, DOI 10.1242/jcs.074849 (2011).

128. Y. Yanase et al., Immunol. Cell Biol. 89, 149–159, DOI 10.1038/icb.2010.67 (2011).

129. V. Niggli, J. Rossy, Int. J. Biochem. Cell Biol. 40, 344–349, DOI 10.1016/j.biocel.2007.02.012 (2008).

130. S. Martinelli et al., Front. Immunol. 4, 84, DOI 10.3389/fimmu.2013.00084 (2013).

131. V. Muralidharan-Chari, J. W. Clancy, A. Sedgwick, C. D’Souza-Schorey, J. Cell Sci. 123, 1603–1611, DOI 10.1242/jcs.064386 (2010).

